# Iron Memory in *E. coli*

**DOI:** 10.1101/2023.05.19.541523

**Authors:** Souvik Bhattacharyya, Nabin Bhattarai, Dylan M. Pfannenstiel, Brady Wilkins, Abhyudai Singh, Rasika M. Harshey

**Affiliations:** Department of Molecular Biosciences and LaMontagne Center for Infectious Diseases, University of Texas at Austin; Austin, TX 78712; Electrical & Computer Engineering, University of Delaware, Newark, DE 19716

**Keywords:** Swarming, Memory, Iron, E. coli, Antibiotics, Biofilms, Time-delay model

## Abstract

The importance of memory in bacterial decision-making is relatively unexplored. We show here that a prior experience of swarming is remembered when *E. coli* encounters a new surface, improving its future swarming efficiency. We conducted >10,000 single-cell swarm assays to discover that cells store memory in the form of cellular iron levels. This memory pre-exists in planktonic cells, but the act of swarming reinforces it. A cell with low iron initiates swarming early and is a better swarmer, while the opposite is true for a cell with high iron. The swarming potential of a mother cell, whether low or high, is passed down to its fourth-generation daughter cells. This memory is naturally lost by the seventh generation, but artificially manipulating iron levels allows it to persist much longer. A mathematical model with a time-delay component faithfully recreates the observed dynamic interconversions between different swarming potentials. We also demonstrate that iron memory can integrate multiple stimuli, impacting other bacterial behaviors such as biofilm formation and antibiotic tolerance.

## Introduction

Constantly changing environments present a challenge to the survival of all organisms^1^. They can adapt to change either by behavioral or by phenotypic plasticity^2^, but must decide among available phenotype choices^3^. Response times during decision making can be critical for survival^3^. Vertebrates use their nervous system for faster decision-making by storing information about their prior experiences^4^. The process of storage and retrieval of information is called memory^4, 5^, which can be stored for a few seconds to several years^6^. A stronger memory can be brought to bear on the system by ‘conditioning’^7^ – a process where repeated encounters to a stimulus can be stably linked to a specific response^8, 9^, drastically reducing decision-making time^10^.

Bacteria experience large environmental fluctuations to which they can readily adapt^11^. They commonly do this by transducing environmental signals to elicit appropriate transcriptional responses^12^ that occur on the time scale of minutes, dissipating in the absence of the signal. Keeping these systems always ’on’ is costly^12^, so bacteria also employ less-costly bet-hedging strategies that stochastically switch among many alternative phenotypic states in similar time frames ^13, 14^. Alternatively, different cell types can pre-exist in an isogenic population and respond heterogeneously to antibiotic stress^15^ or to starvation^16^. All of these strategies come under the umbrella of bacterial memory^17^.

Whether bacteria can store memory has been explored both theoretically^17, 18^ and experimentally^19–22^. Memory can influence the fitness of individuals^18^ or of a community^19^, as well as bacterial interaction with their host^20^ and defense against phages^23^. The mechanism of memory storage in bacteria involves genetic^5^ or epigenetic factors^24, 25^ such as type 1 fimbriae phase variation^5^, or motile-sessile transition^26^ or chemotaxis^27–29^. These varied molecular mechanisms of bacterial memory storage are more stimulus-specific in nature, in contrast to the nervous system where no matter the type of stimulus, the same molecular principle of storage is employed^4^. The existence of the latter kind of memory mechanism in bacteria can only be conjectured from some recent reports^21^ ^22^.

In this study, using *E. coli* swarms as our experimental system, we asked a question not considered before: is there a heritable memory in bacteria that can integrate multiple stimuli? The choice of this system was based on observations that a prior experience of swarming hastens the onset of swarming. Swarming motility is a collective, flagella-driven, adaptation to colonizing the surface of semi-solid media by population expansion^30–34^. *E. coli* will typically swarm on semi-solid media (∼0.5% agar), but are non-motile on solid or hard media (∼1.5% agar). The ’softer’ agar is more conducive to swarming because among other properties, it also holds more water, allowing flagella to work on a surface terrain. Surface colonization presents varied challenges, both physical and nutritional^33^. Nutritional challenges include acquisition of iron^35^ leading to upregulation of efflux pumps essential for the secretion of iron-binding siderophores into the extracellular matrix, as well as upregulation of iron import systems^36–40^. When bacteria cultured in liquid media are transferred to swarm media, a long lag ensues before swarming begins^32^. Widespread transcriptome changes accompany the onset of swarming^38, 41–44^, revealing that the bacteria are adapting to a distinctly different environment during the lag phase. If, however, bacteria from a swarm are transferred to a fresh swarm media, the lag is considerably shortened^45^, suggesting that the bacteria may be already ’conditioned’ to swarm.

To interrogate whether swarming bacteria ’remember’ the adaptation mechanisms employed while swarming, we designed an experimental set-up that allowed monitoring of swarms initiated from >10,000 single cells. We report here that the bacteria indeed possess a heritable memory of the swarming state that is retained on the time scale of hours over at least four generations. We show that the mechanism of this type of memory storage is primarily related to how much iron is stored intracellularly. Other cues that promote biofilm formation and antibiotic tolerance are also routed through iron memory, reflecting the essential role iron plays in central metabolism. Our findings highlight a more general physiological mechanism of memory storage in *E. coli*, which we predict will be widely present in other bacteria.

## Results

### Swarming proficiency pre-exists in planktonic cells

The impetus for experiments performed is this study originated from a serendipitous observation that swarm cells plated for determining CFU counts (colony forming units) on freshly-made (hence moist) hard agar, produced colonies whose phenotype was different from those made by planktonic cells (growing in liquid culture) plated on similar agar. Compared to planktonic (P) colonies, swarm-derived (S) colonies had more irregular margins, were larger, and flatter (when viewed from the side) (Fig. 1, a-b). We posited that these differences stemmed from prior ’conditioning’ of S cells, so they were attempting to swarm even on hard agar because the plates were still moist. We quantified these morphologies by measuring colony circularity (using both area and perimeter), and size (diameter) (Fig. 1c); S colonies were significantly less circular and larger. On closer inspection, a fraction (∼20%) of P-derived colonies also displayed S-colony characteristics, although less prominently (Fig. 1b, top; compare #9-10 with #1-8), suggesting that P-cells were also heterogeneous for swarming proficiency, with some inherently primed for swarming. To test the latter deduction, we performed Single Cell Inoculum (SCI) swarm assays, where P cell-derived swarms were initiated from single P cells isolated using the dilution-to-extinction method^46^ targeted to yield an average of 0.5 cells per unit volume (Fig. 1d and Extended Data Fig. 1, a-b). The sampling events would follow a Poisson distribution, so ∼40% of seeded swarm plates would have received a cell. This was confirmed by parallel growth in microtiter plates (Extended Data Fig. 1c, top), which matched the expected positive sampling events with a high degree of precision (Extended Data Fig. 1d). Therefore, in all SCI assays, at least 120 swarm plates were inoculated to gather data for 50 swarms (Extended Data Fig. 1c, bottom), again with high precision (Extended Data Fig. 1e). The sample size was kept sufficiently large to increase confidence in the statistical analysis. Additionally, such Poisson distributions would sample multiple cells in ∼5% of events. We could identify those events on a swarm plate by just counting the number of swarm ’nucleation’ centers and exclude them from our analysis (Fig. 1e). In parallel, we compared data from SCIs versus 100-or 10000-cell inoculums, where any heterogeneity present in single cells is expected to be subsumed within the large cohort. Swarming proficiency was calculated as distance covered over time (Fig. 1f). SCIs showed a large variability in swarming, in contrast to the larger cell inoculums (see the endpoints of lines in each group). The comparison of this variability between the three data sets was complicated by the fact that each cohort had a different lag period, as expected from the dependence of lag on cell-density^33, 47^. A zone-adjustment was therefore performed by arbitrarily dividing the plate into 3 zones (C/M/O) (Fig. 1g) and comparing swarm diameters of every cohort within each zone (Fig. 1h; see Methods). The zone-wise variability in frequency distributions (FD) of swarm diameters (5 mm bins) were then compared (Fig. 1, i-k), and the noise calculated (Fig. 1, l-n). In every zone, SCIs had significantly higher levels of noise (a reflection of heterogeneity) compared to other cohorts, the maximal noise evident in the outermost zone O (Fig. 1k and n). The large heterogeneity observed within the P cell-derived SCIs reveals that swarming proficiency pre-exists even in planktonic cells.

**Fig. 1.**
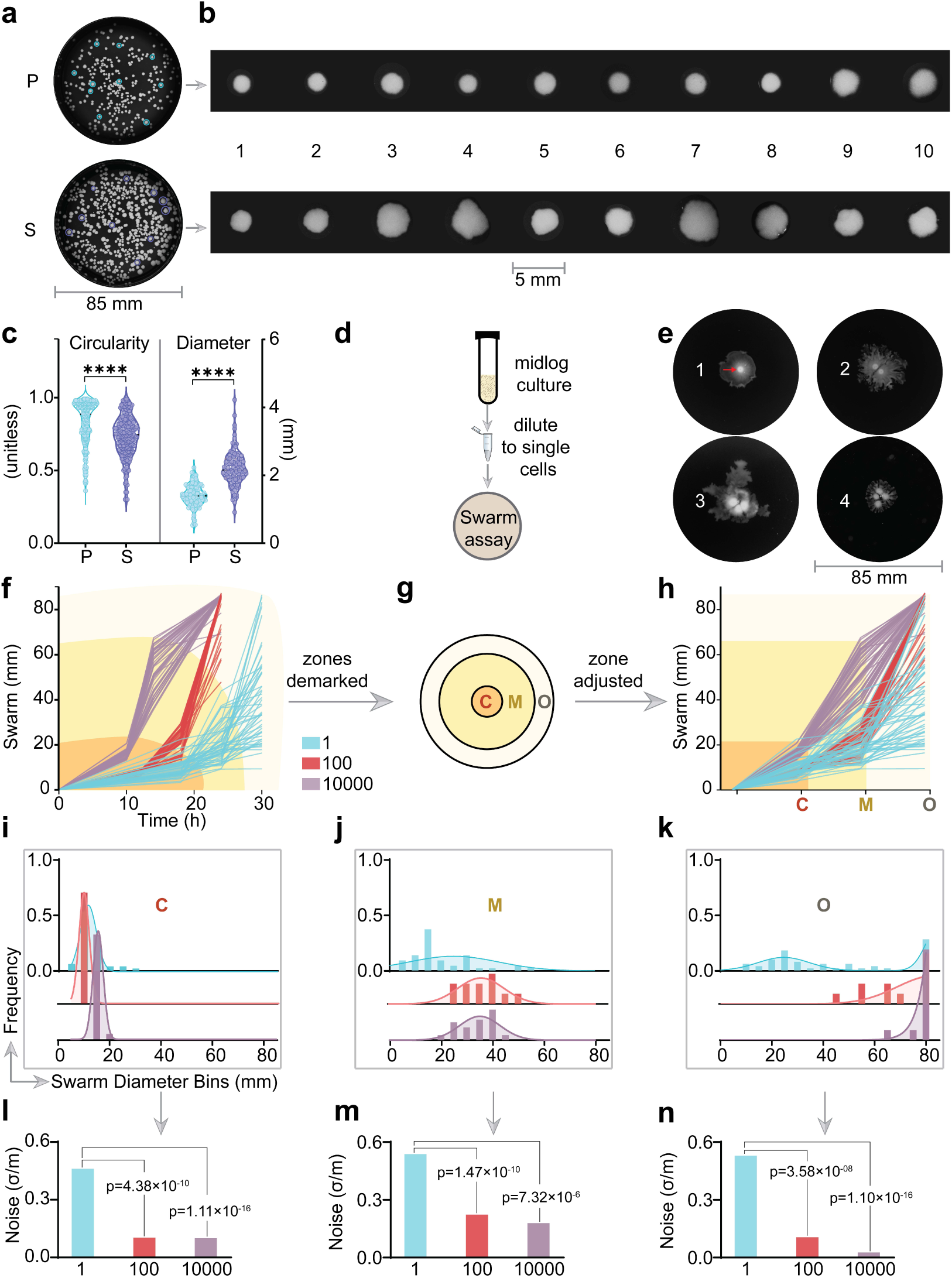
Swarming proficiency pre-exists in planktonic cells. **a**, Representative hard-agar plates showing CFUs of planktonic (P) cells from a mid-log phase liquid culture, and swarm (S) cells collected after growth for 16h. In both cases, a 10^-6^ dilution of 0.5 OD cells was plated on hard agar plates and incubated for 16h at 37°C. **b**, Blown-up images of 10 randomly selected colonies from A (indicated by colored circles). **c**, Distribution of physical parameters of P and S colonies (n=218). See Methods for circularity calculations. [****p <0.0001, Mann-Whitney test] **d**, Flow chart of single P-cell swarm assays, where the dilution was estimated to yield a single cell in a unit volume of 4ul (see Extended Data Fig.1 and Methods for details). **e,** Swarms with varying number (1-4) of swarm nucleation centers (red arrow) dictated by the initial number of cells spotted on the plate. **f**, a spaghetti plot showing swarm diameters (Y-axis) measured over time (X-axis). Each line represents a replicate. **g,** Three arbitrary zones considered for adjustment: C, central, M, medial; O, outer. **h,** zone-adjusted comparisons (see Methods and text). **i-k**, Frequency distribution of swarm diameters (5 mm bins), color-coded as in (**f**). **l-n**, Comparison of noise (SD/mean) within cohorts. p values were calculated from F-tests (see Methods).

### Swarming-related memory is present in both P and S cells

To explore whether the heterogeneity of swarming proficiency was random or inherited, we evaluated the swarming capability of offspring from single cells over several generations (Fig. 2a). The starting cells represent our G0 generation ’mothers’, and were derived from both P and S conditions. We first standardized the established dilution-based protocol^46^ to precisely collect daughter cells at different generations (Extended Data Fig. 2, a-c). Some G0 mothers (n=50) were directly spotted on swarm plates (Fig. 2, b-c and Extended Data Fig. 2d). Others were grown in 96-well plates for either 4 (Fig. 2, d-e) or 7 (Fig. 2, f-g) generations (n=25, x axis) before plating single daughters of each mother (16 expected, Y axis in Fig. 2, d & f) on swarm media (see Extended Data Fig. 2, d & f). We initially probed other generations as well; but to reduce time and effort in these labor-intensive experiments, we confined our observations to these generations, which gave conclusive results during our initial screening. In the bubble plots (Fig. 2, b-g), the size of each bubble represents the swarm diameter and the color represents the noise within 16 daughters of the same mother. The behavioral patterns of P mothers (PèS data, b-d-f) was distinctly different from the S mothers (SèS data, c-e-g). P mothers (G0) showed the expected heterogeneity (see 1-cell data in Fig. 1f) in swarming (Fig. 2b, Supplementary Fig. 1). In contrast, all the G4 daughters descended from a single P mother were more homogeneous (Fig. 2d, follow the data vertically or column-wise for the 16 siblings). Some mothers (#11, #13, #14) showed variability, which could have come from experimental noise. A K-means clustering of the means derived from each column resulted in three distinct groupings labeled XS (poor), M (moderate) and L (efficient) swarmers. The majority patterns, i.e., consistent behavior of each sibling group, suggested that some information stored in the mother was passed down to the daughters. This information was lost by G7 as the variability returned (Fig. 2f, Supplementary Fig. 1). In summary, P daughters inherit some form of memory that is intact up to four generations, but is lost by the seventh generation, restoring the original heterogeneity.

**Fig. 2.**
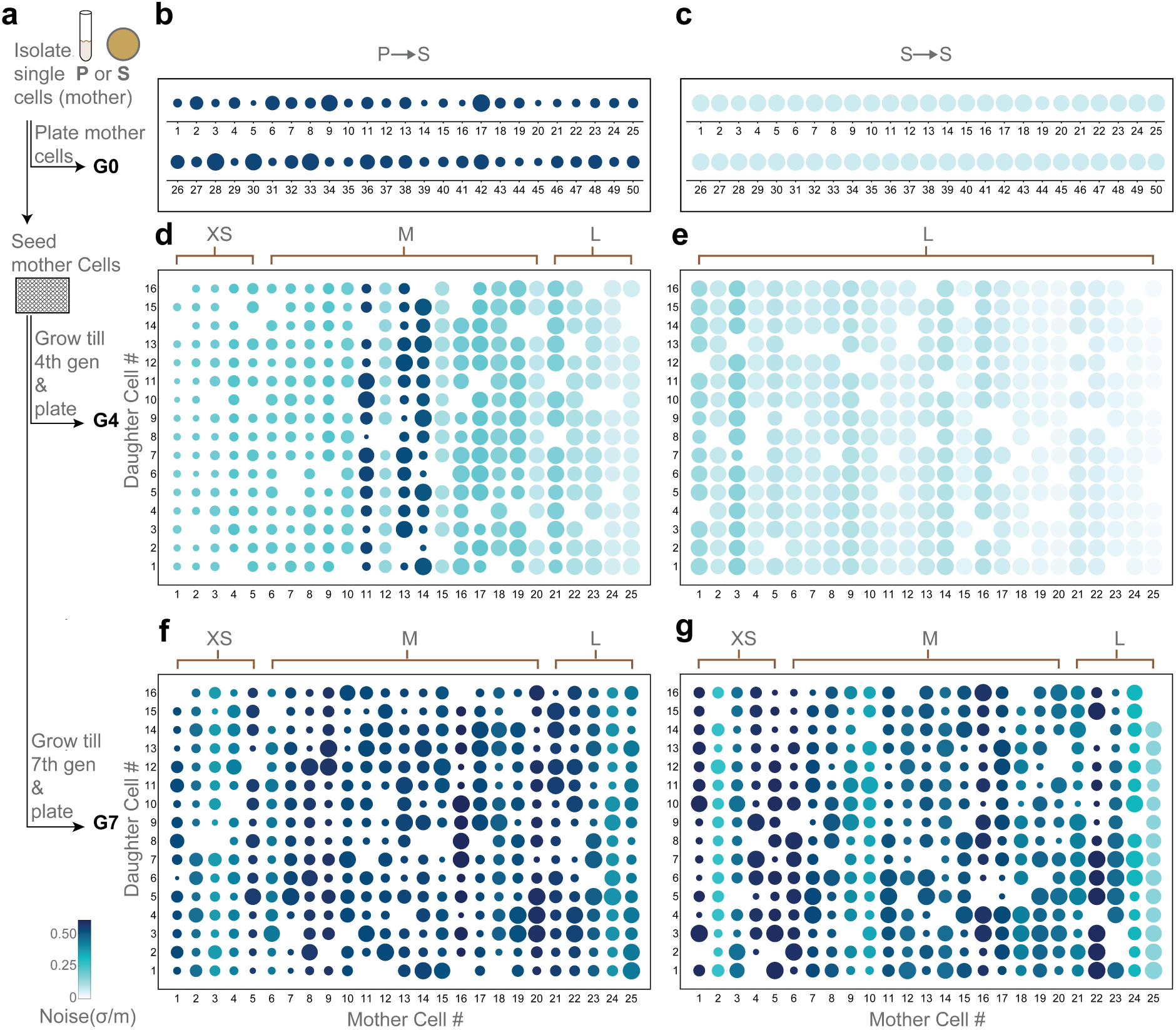
Swarming-related Memory is present in both P and S cells. **a,** Flowchart summarizing the operations involved. P and S cells were collected and diluted to get single cells; these are the starter G0 mother cells which were directly spotted on swarm plates (PèS), as well as seeded in 96-well plates and grown for either 4 or 7 generations (see Extended Data Fig.2 for the standardization process of the generations and statistics). All the resulting daughter cells (16 from each mother) were further similarly diluted and spotted separately on swarm plates. **b-g**, Bubble-array plots where each bubble represents swarm diameters from SCIs (∼1700 total plates with swarms). The size of the bubble represents the swarm diameter and the color represents the noise within a group of daughters born from the same mother (**b**-**c:** G0; **d**-**e:** G4; **f**-**g**: G7; **b**, **c**, **d**: P mother; **e**, **f**, **g:** S mother). See Supplementary Fig. 1 for the noise data of each mother. Based on the daughters’ ability to swarm in G4 and G7, a K-means clustering of mother cells resulted in three distinct groups; XS – poor, M – moderate, and L – efficient swarmers.

The same set of experiments performed on S mothers (SèS data, c-e-g) showed a very different outcome at G0 and G4. In contrast to P mothers, swarm diameters of S mothers at G0 were uniform (Fig. 2c, Supplementary Fig. 1), suggesting that they retained ’swarm’ memory or were ’conditioned’ in some way while swarming. Similar to P cells, this memory was maintained in S daughters until G4, but with even lower noise (Fig. 2e). In addition, the K-means clustering resulted in only in one group (L), in contrast to 3 groups in P daughters. By G7, the S daughters had lost this memory, similar to P cells (Fig. 2g, Supplementary Fig. 1), but even here the noise was lower than in P cells. In summary, the uniform behavior of G0 S mothers, as well as the single cluster of G4 daughters indicates that S cells are conditioned, and that this swarm memory is retained at least for 4 generations but lost by the 7th generation.

In conclusion both P and S cells have a heritable multigenerational memory of swarming, henceforth referred to simply as memory. While this memory is heterogeneously distributed in P cells, it is homogenously present in S cells.

### Cells have an iron memory

What is the nature of this multigenerational memory? Given the significant variation in the swarming proficiency of SCIs, we reasoned that any factor(s) that alters this heterogeneity would provide insights into its nature. Since swarming involves both growth and motility, we first looked at the SCI variation in these two factors. We measured the total pixel intensity of each swarm as the growth variable (V_G_) and the maximum swarm diameter as the motility variable (V_M_) (Extended Data Fig. 3a). A K-means clustering (Extended Data Fig. 3b) of the V_G_-V_M_ data from P mothers at G0 resulted in four distinct groups – poor growth & motility (I), moderate growth & motility (II), moderate growth & good motility (III), and good growth & moderate motility (IV). The actual plate images from these clusters revealed that the clustering was real and could be visually correlated (Extended Data Fig. 3c). Cohorts that showed less variation (G4 P, G0 S, 100-cell,10000-cell) exhibited only two clusters (Extended Data Fig. 3b) but could not be visually correlated (Extended Data Fig. 3c), suggesting the presence of only a single cluster in these samples. Interestingly, the four real clusters of G0 P cells that disappeared at G4, reappeared in G7 (Extended Data Fig. 3c, 1P-G7), consistent with memory retention for 4 generations but not for 7. Together, the clustering analysis confirmed that V_G_ and V_M_ are both important for swarming heterogeneity.

**Fig. 3.**
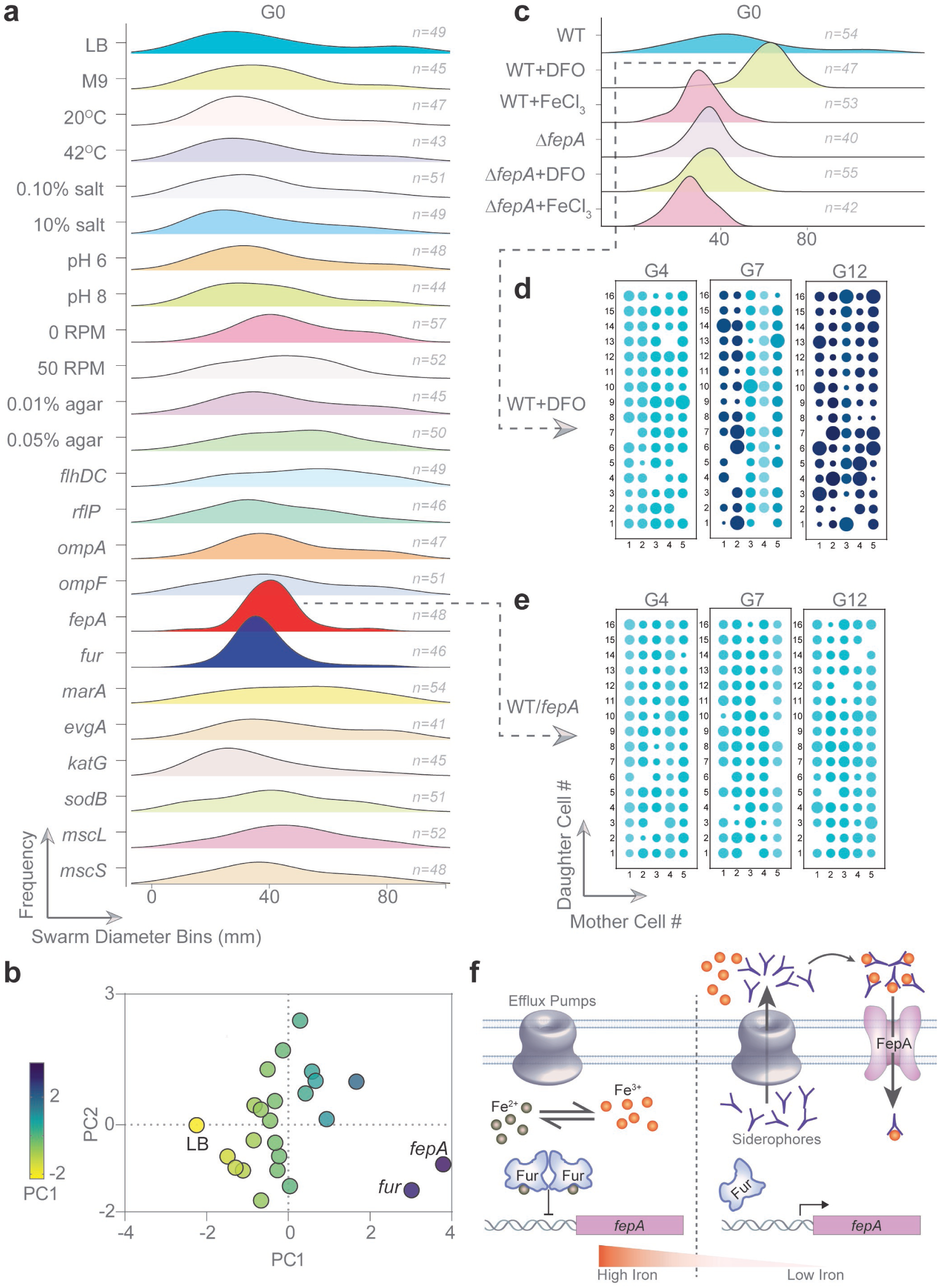
Cells have an iron memory. **a**, Ridgeline plots showing frequency distribution (0 to 1 in each plot) of swarm diameters of G0 P cell-SCIs (∼2400 total), where prior to plating, P mothers were either subjected to indicated growth environments (in liquid), or transformed with plasmids expressing genes known to influence swarming. See Supplementary Fig. 2a for the noise data. **b**, Principal Component Analysis of data from **a**. The mean, SD, and noise were considered as variables of each condition. Eigenvalues of PC1 (60.06%) and PC2(39.87%) are expressed on the X-and Y-axis, respectively. Each dot represents a condition with relevant ones labeled. See Supplementary Fig. 2b for the biplot. **c**, Ridgeline plots specific to WT or *ΔfepA* P cells experiencing iron-poor (DFO) or iron-rich (FeCl3) conditions. **d-e**, bubble-array plot of G4, G7, and G12 daughter P cell swarm assays (as in Fig. 2) of WT strains either starved for iron (DFO) or had abundant iron. **f**, Schematic of Iron homeostasis as controlled by Fur, the global negative regulator of intracellular iron. See text for details.

To explore environmental or genetic factors that might reduce the swarming heterogeneity of the P mother cells (measured as swarm diameters), we perturbed several factors prior to inoculation of these G0 cells on swarm plates. The environment was perturbed by changing several parameters such as media, temperature, pH etc. (Fig. 3a, Environmental; see Supplementary Fig. 2a for noise data), but no apparent decrease in swarming heterogeneity was observed. These observations were further quantified by performing a Principal Component Analysis (PCA). Since the mean and the noise are major descriptors of sample data, these values were taken as variables for PCA^48^. None of the environmental perturbations changed principal components significantly (Fig. 3b, Supplementary Fig. 2b). Genetic factors were assessed by introducing plasmids carrying representative genes that had been identified as contributing to *E. coli* swarm physiology^38, 44, 49, 50^. These genes encoded for: porins (*ompA, ompF*), iron regulation (*fepA, fur*), efflux (*evgA, marA*), redox (*katG, sodB*), and mechanosensing (*mscL, mscS*). Of these, perturbation of iron homeostasis significantly reduced noise and clustered the data very differently in PCA when compared to LB control (Fig. 3b, *fepA* and *fur*; 3b; Supplementary Fig. 2b). Iron starvation is known to be crucial signal for swarming^37, 44^. FepA is an outer membrane transporter for ferric enterobactin, while Fur represses iron uptake genes^51^. Compared to WT, Δ*fepA* and Δ*fur* strains showed no significant differences in growth or swimming motility as assayed in soft agar (0.3% w/v) (Extended data Fig. 4, a-b), yet had different swarming phenotypes: the Δ*fepA* strain was a poor swarmer, whereas Δ*fur* did not swarm (Extended data Fig. 4, c-d). The importance of iron in maintaining swarming heterogeneity was confirmed by growing P mothers in iron-rich (FeCl_3_) or iron-starved conditions (deferoxamine mesylate or DFO, an iron chelator) before conducting SCI assays (Fig. 3c,). These treatments severely reduced the heterogeneity, creating defined peaks (Fig. 3c). A shift to the right on the graph indicates better swarming (+DFO), while that to the left indicates the opposite (+FeCl_3_), validating studies showing that iron starvation is a swarming signal^37, 50^. Interestingly, neither DFO nor FeCl3 had any effect on Δ*fepA* mothers (Fig. 3c), consistent with the fact that iron uptake is impaired in this mutant.

**Fig. 4.**
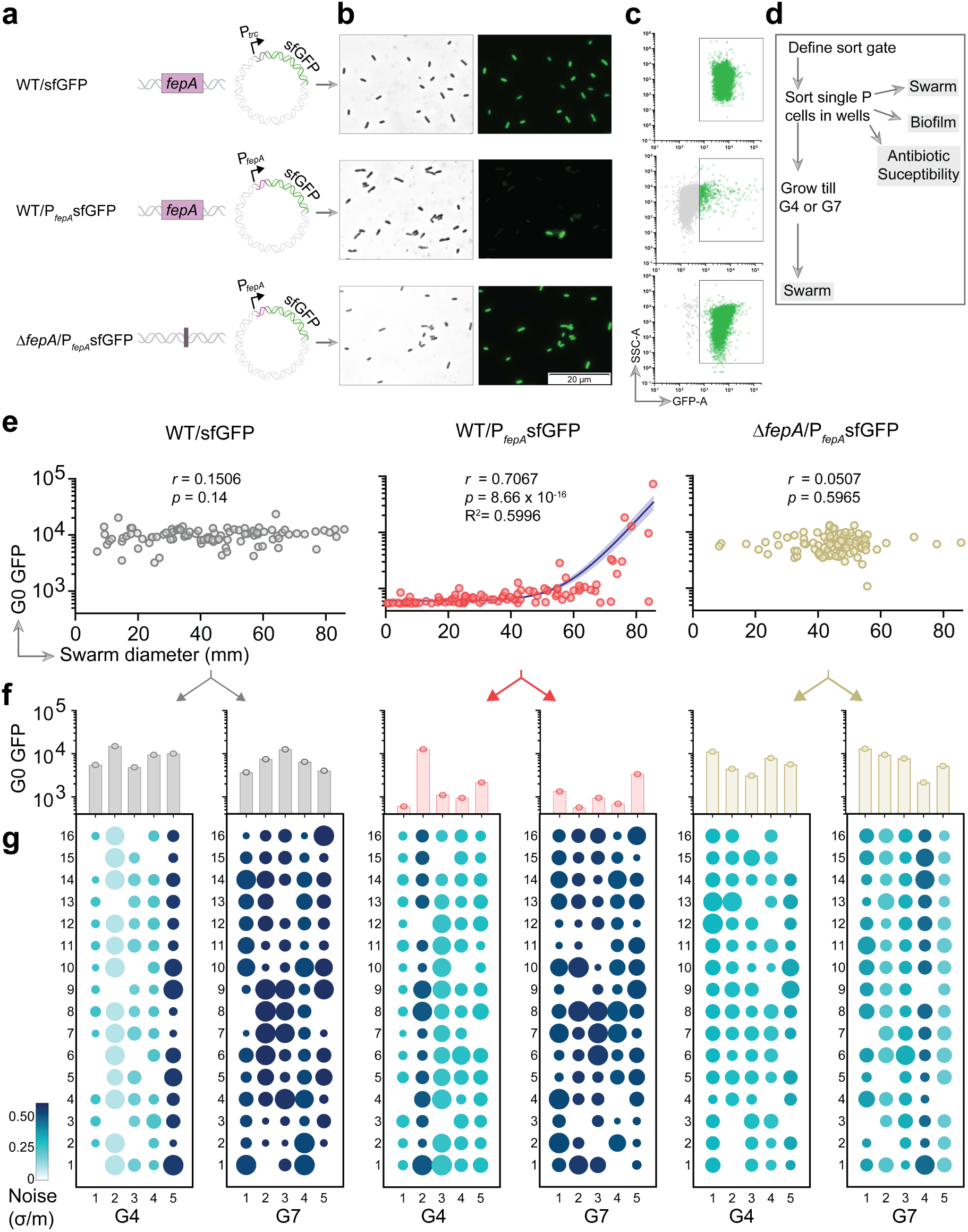
Swarming iron memory tracks from mother to daughters. **a.** WT or Δ*fepA* cells harboring Ptrc-sfGFP or P*fepA-*sfGFP plasmids_._ See Supplementary Fig. 3 for cloning details. **b**, Brightfield (left) and fluorescence (right) microscopic images of strains depicted in **a**. **c**, Results of FACS analysis of strains indicated in **a**. Singlets were gated based on SSC-A (side scatter channel -area) and GFP intensity. A sort gate was defined to collect singlets where the background fluorescence of WT cells without GFP was defined as the minimum threshold See Extended Data Fig.5 and Methods for details). Gray, cells below threshold. Green, cells within the defined gate. **d**, Flow chart of how sorted cells were used. **e**, Correlation of single-cell GFP intensities (from FACS, n=96) with swarming of G0 mother cells collected from the different strains indicated. r, spearman correlation coefficient, p value as indicated. The blue line represents the exponential regression line with indicated R^2^ values. The blue error band represents 95% confidence interval (CI). **f**, Fluorescence intensities of sorted G0 mothers that were dropped into one well of a 96-well plate, indicated as bars. **g**, The G4 or G7 daughters of the mothers with known GFP intensities (indicated as bars in **f**) were assayed for their swarming ability and the results are shown as bubble-matrix plots. See Supplementary Fig. 4 for the FACS data of each experiment.

To investigate the effect of iron starvation on memory, we followed memory inheritance for G4, G7 and G12 daughters in WT P cells treated with DFO. Memory was now largely maintained till G7 (Fig. 3d), but was lost by G12, showing that experimentally induced iron starvation extends memory beyond G4. This suggests that memory might persist for longer durations by increasing intracellular iron. Overexpression of *fepA* (expected to import iron) indeed retained memory till G12, the last generation we tested (Fig. 3e).

In summary, intracellular levels of iron are an agent of memory that can be either environmentally or genetically conditioned.

### Swarming iron memory tracks from mother to daughters

Most methods for measuring intracellular iron require complex endpoint chemistry^52^ and those that measure iron in live cells can impact cellular physiologies^53^. To avoid such experimentally-induced perturbations while quantifying iron in our setup, we designed a fluorescent reporter-based iron biosensor (Supplementary Fig. 3). The reporter is sfGFP, transcribed from the Fur-regulated *E. coli fepA* promoter. Fur is the global repressor of iron uptake^35^ (Fig. 3f). The *fepA* promoter was used to drive sfGFP because this gene has a strong Fur binding site (see Table I in^54^), is crucial for swarming^55^, and is highly expressed during swarming^44^. When intracellular iron levels are high, Fur stops further iron intake by complexing with iron and binding to a ‘Fur box’ adjacent to promoters of hundreds of iron homeostasis genes^56^ to repress transcription (Fig. 3f). When iron levels drop below a threshold, the system is depressed and genes involved in siderophore production, secretion (efflux pumps), and uptake (siderophore receptors) are expressed (Fig. 3f). Thus, high intracellular iron should cause strong suppression by Fur yielding a low fluorescence in our setup and vice versa. The sfGFP fluorophore was expected to yield a signal strong enough^57^ to be detected even under strong Fur repression. A control plasmid expressing sfGFP from a P_trc_ promoter in WT cells gave a homogeneous background signal (Fig. 4a-b, top), whereas sfGFP driven by the *fepA* promoter (P*_fepA_*) produced a heterogeneous signal, reflecting heterogeneity of iron levels (Fig. 4a-b, middle). The ability of the biosensor to report on cellular iron levels was further tested by subjecting WT/P*_fepA_*sfGFP cells to FeCl_3_ and DFO treatment. Not only did fluorescence decrease and increase, respectively, the heterogeneity in signals also reduced, as expected (Extended Data Figure 4e; heterogeneity due to different plasmid copy numbers is controlled for by the IPTG-induced, continuously ’ON’ sfGFP plasmid). When P*_fepA_*sfGFP was introduced into the Δ*fepA* strain, which cannot import iron, the fluorescence signal became homogeneous, as expected if all cells had low iron (Fig. 4a-b, bottom). FACS analysis confirmed results from microscopy (Fig. 4c, Extended data Fig. 5): only WT/ P*_fepA_-*sfGFP showed heterogeneity (Fig. 4c, middle, green singlets). Having validated this construct as an iron biosensor, we used it next to validate iron memory in swarming, as well as other phenotypes expected to be impacted by iron levels (Fig. 4d).

**Fig. 5.**
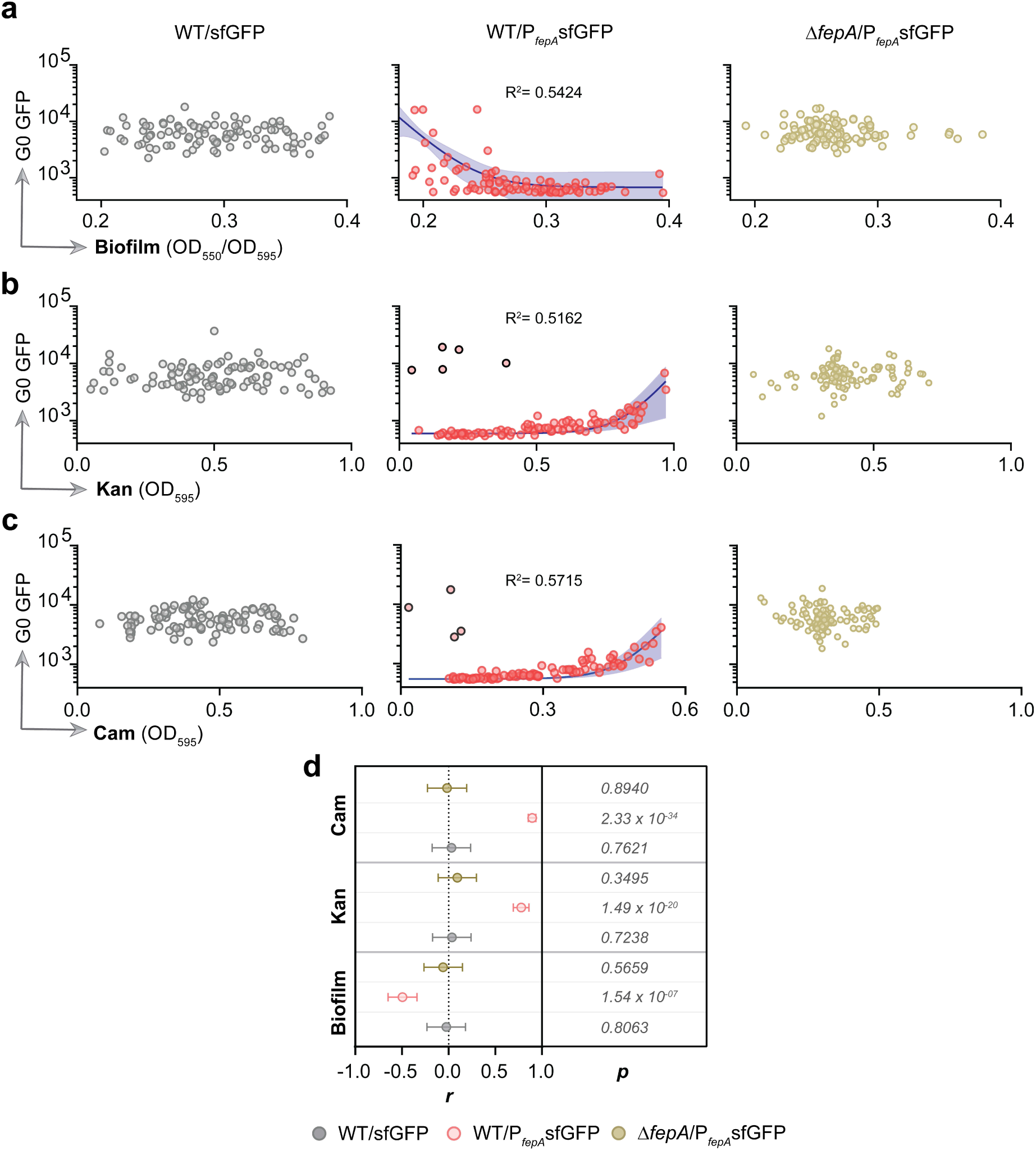
Iron memory impacts biofilm formation and antibiotic survival. **a-c**, Correlation of single-cell GFP intensities of G0 mother cells from FACS (n=96, for each sample) with: **a,** biofilm formation; **b-c**, antibiotic survival (2 μg/ml Kan & 1.5 μg/ml Cam). Blue lines represent exponential regression lines with indicated R^2^ values. The blue error bands represent 95% CI. **d**, Distribution of r, the spearman correlation coefficients of GFP intensity and growth under the various conditions tested. The error bars represent 95% CI. *p* values are given on the right. See Supplementary Fig. 4 for the FACS data of each experiment.

FACS was used to first sort P cells harboring all the biosensor-harboring constructs through a pre-defined sort gate that included a range of signals covering low to high iron levels (Extended Data Fig. 5 and Supplementary Fig. 4). Single cells covering this range were then dropped on swarm plates to measure swarming proficiency (Fig. 4d). Although cells with the control P_trc_ plasmid showed heterogeneity in swarming (Fig. 4e, left, follow x-axis), no correlation between the sfGFP signal and swarming diameter was observed (Fig. 4e, left). As expected, however, a positive correlation between these two measurements was observed in WT/ P*_fepA_* sfGFP G0 cells, i.e., cells with high sfGFP (low iron) were better swarmers (Fig. 4e, middle). In Δ*fepA/* P*_fepA_*-sfGFP strain, where iron import is impaired, we observed a clustering of swarm diameters (Fig. 4e, right, follow x-axis), consistent with the reduction of heterogeneity (see Fig. 3b, Δ*fepA*). Next, for each of the strains shown in Figure 4e, we performed a multigeneration analysis with randomly sorted G0 cells (see Supplementary Fig. 4 for sort results). Five such random P mothers from each strain were propagated until G4 or G7 (Fig. 4f, g). Unlike in Fig. 2a, where the physiological state of the seeded mothers was unknown, this time we had knowledge of their iron levels based on their fluorescence levels, shown as bars (Fig 4f, y axis - log scales; high bars = low iron). The swarming proficiencies of each sibling set is plotted as before (follow the columns below each bar for daughter cell data). The swarm diameters of daughters correlated closely with iron levels of the mother. For example, in the G4 cohort of WT/ P*_fepA_*sfGFP mothers (Fig 4g, third panel from the left), mother #1 (first column, short bar=high iron) birthed daughters that were all poor swarmers, while mother #3 (third column, higher bar=low iron) birthed daughters that were efficient swarmers. The G7 data from these mothers mirrored previous observations that iron memory is lost by G7 (Fig. 4g, fourth panel). In contrast, no such correlation between iron and swarm diameters was observed in the other two strains shown in Fig. 4e (left, right). In both strains, memory was maintained till G4 (Fig. 4g, first and fifth panel from the left). The Δ*fepA* strain exhibited high sfGFP signals due to their low intracellular iron (Fig 4e, right, follow y-axis). Interestingly, memory of low iron in this Δ*fepA* strain was maintained at least till G7, which is an improvement upon the G4 memory of WT cells (Fig. 4g, second and sixth panels from the left). This observation, along with the data in Fig. 3e, revealed that both decreases or increases in iron levels can extend the span of memory beyond WT levels, although the phenotypes in both cases are at the opposite ends of the spectrum.

In summary, we constructed and validated an iron biosensor that can be used to quantify the iron memory and sort cells based on the quantification. Results from this biosensor showed a strong correlation of iron memory with swarming, which is propagated from mothers to daughters till G4 in WT cells.

### Iron memory impacts biofilm formation and antibiotic survival

Besides swarming, iron is also crucial for several physiologies^58^. We tested a few such phenotypes for their correlations with intracellular iron levels (Fig. 5, a-d, Supplementary Fig. 4). Recent studies have shown that iron is crucial for the decision to enter the biofilm state^59, 60^, which is a contrasting sedentary lifestyle choice when compared to swarming^34^. To test if iron memory is a predictor of biofilm formation, we measured this ability in G0 P cell SCIs using a standard crystal violet staining assay^61^ (Fig. 5a). As expected, we observed no correlation between iron and biofilm formation in the control WT/P_trc_-sfGFP strain (Fig 5a, left; see 5d for correlation data), but a strong anticorrelation between low iron and biofilms in WT/ P_fepA_ sfGFP cells (Fig. 5a, middle; 5d), i.e., more iron is a signal for biofilms. This anticorrelation was absent in strains lacking FepA (Fig. 5a, right; 5d). These data are consistent with the opposing ’move vs stay’ decisions associated with the two behaviors.

Iron is critical for redox homeostasis^35, 62^, which is upended by antibiotics^63, 64^. To test if iron levels can be a predictor for antibiotic survival, sorted cells of the same three strains shown in Fig. 5a were treated with ∼½ MIC of the antibiotics Kanamycin (Kan) and Chloramphenicol (Cam) for 4h. A positive correlation was observed between low iron and increased survival with both antibiotics (Fig. 5b-d, Supplementary Fig. 4). This correlation was absent in Δ*fepA/* P*_fepA_*-sfGFP or P_trc_-sfGFP strains.

In summary, iron memory is not restricted to swarming decisions alone, but plays a crucial role in several bacterial physiologies.

### A mathematical model that explains the propagation and switching of iron memory states

To understand how iron memory can be propagated from mother to daughter, we thought it prudent to first understand the phenomenon mathematically, before pursuing detailed future investigations into underlying molecular mechanisms.

Given that cells switched dynamically over multiple generations among three different states (XS, M, L; Fig. 2), we used an ordinary differential equation model as our mathematical framework (see Methods). The model assumes different rates of switching between the states (Fig. 6a). Starting from any state, cells could stochastically switch to other states over time, ultimately recreating the original phenotypic heterogeneity (Fig. 6b-d). In order to satisfy the curious observation that the S data had lower noise, i.e. S daughters took longer to switch compared to P daughters (compare Fig. 2d to 2e and 2f to 2g), a time delay component had to be incorporated. Model parameters such as the delay duration and kinetic rates were inferred by fitting the model to data at G4 (Fig. 2d-e), and these rates were further constrained so as to recreate the intercellular variation seen in single G0 mothers by G7 (see Methods). Our modeling results reveal that in the case of high to moderate iron states (XS and M), all three states started to appear during G3-G4 transition (Fig. 6b-c). In contrast, the low iron state cells (L) initiated this process at G4-G5 transition suggesting that conditioning (prior swarming experience) can enforce iron memory further than stochastic switching (Fig. 6d). Our model could capture the real data efficiently as seen when comparing fractions of different cell states in Figs. 6b-d (dashed lines for G0 data) to the experimental data in Fig. 2 (see Supplementary Fig. 5). In other words, no matter the starting cell state, the emerging population eventually converges to the original ratio of cell states. Using the model data, a cell lineage dendrogram was generated, which visually illustrates how these cell states can reappear (Figs. 6-e-g).

**Fig. 6.**
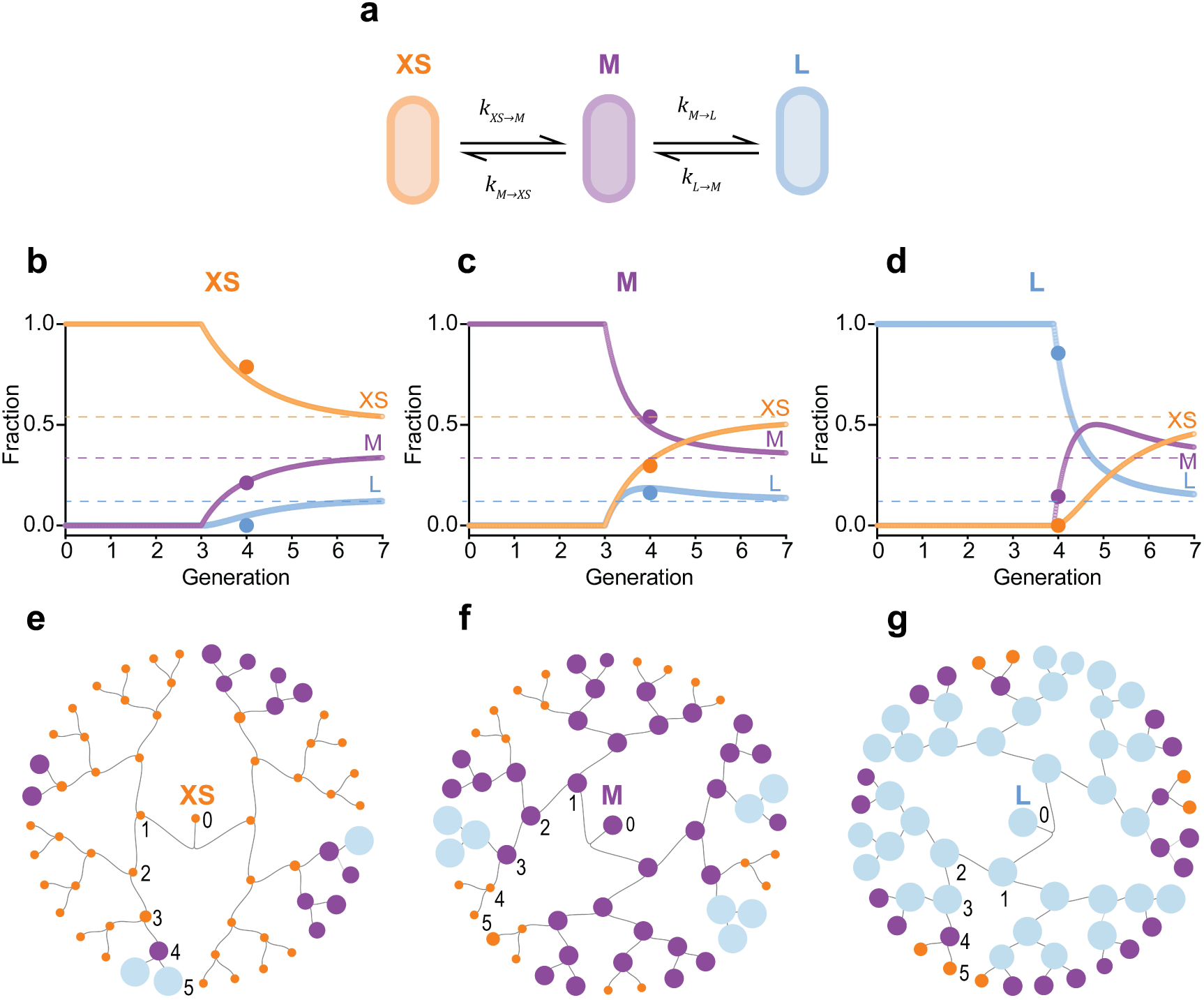
A mathematical model that explains the propagation and switching of iron memory states. **a**, Depiction of the mathematical framework used to model the data in Figure 2. Briefly, XS, M, and L cells with poor, moderate, and efficient swarming capabilities respectively can switch to other states by the given rates with a time delay. See text and Methods for details. **b-d**, Results from the time delay model showing how all three swarming states can be spawned starting from any cell type (as indicated on top). The XS and M cells can maintain the memory till G3 and then start to switch to other cell types, whereas L cells take until G4 before switching. The dashed lines show the fractions of cell types obtained from G0 mothers data of Fig. 2b (also see Supplementary Fig 5). Irrespective of the initial cell type, the resulting population converges to the original ratio of the cell types. **e-g**, A cell lineage illustrating the memory model. Starting from a specific cell type, the fractions of all three cell types in each of the G1-G5 generations (1-5) were obtained from the model in **b-d** to get the approximate number of each cell type. That information was used to create the circular dendrograms.

In summary, our mathematical model shows that specific rates of dissipation and acquisition of memory can be correlated with the experimentally obtained fractions of cells in different iron states – thus linking the two together.

## Discussion

This study has uncovered a new role of iron in bacterial behavior by demonstrating the existence of physiologically stored ’iron’ memory in *E. coli* that is retrieved over multiple generations. In addition, this memory can integrate multiple stimuli, and thus influence multiple physiologies. To our knowledge, this kind of memory has not been unearthed before, but there is no reason to think that it is limited to the iron pathway monitored in this study.

### Iron memory in swarming

The original observation that led to the present study was a particular colony phenotype present uniformly in swarm (S) cells plated on hard agar, but present non-uniformly in planktonic (P) cells (Fig.1, a-c), suggesting that some attribute of S cells was pre-present in P cells. The latter deduction was confirmed by assaying the swarming potential of a large number of single P cells (Fig. 1, f-n). To test whether the heterogeneity in swarming potential of these cells was random or inherited, the offspring of G0 P cells tracked over several generations were found to inherit their starting swarming potential until G4 but not up to G7 (Fig. 2, b-d-f). The swarming proficiency of S mothers was homogeneous, and also inherited until G4 and lost by G7 (Fig. 2, c-e-g). Thus, both P and S cells have a heritable multigenerational memory of swarming.

Based on prior knowledge of gene regulation during swarming, we tested various candidates that might contribute to swarming memory (Fig. 3a, Genetic). Of these, only the over-expression of critical players in the iron acquisition pathway - *fepA* and *fur* - reduced the noise in G0 mothers significantly. Efflux pumps also play a role in this pathway because they are responsible for siderophore secretion^65^. However, these pumps were not observed to contribute to memory (Fig. 3a, *marA* and *evgA*), ruling out the possibility that the heterogeneity of swarming is caused by the known heterogeneity in efflux pumps^66^. The role of intracellular iron in encoding swarming memory was ascertained using an iron biosensor (Fig. 4). Swarming proficiency tracked with intracellular iron levels, i.e. cells with low iron were better swarmers and vice versa. Memory retention could be manipulated by decreases or increasing iron levels (Fig. 3, d-e), establishing a strong correlation of iron memory with swarming.

### A time delay model for switching of iron memory states

What explains the mixed nature of swarm phenotypes and their heritability (XS, M, and L in Fig. 2)? The short duration of this heritability excludes epigenetic mechanisms^67, 68^, or bistable^69^ and toggle switches^70^. Although epigenetics might play a role in *fur* regulon^71^, a stochastic fluctuation model^72, 73^ could explain the observed fluctuations within the population (Figs. 6a-d). Stochasticity may arise due to intrinsic or extrinsic noise in gene expression^74, 75^. The stochastic fluctuations in transcription and translation of a given gene constitute intrinsic noise (e.g., Fig. 3a, genetic), whereas the effect of other cellular components on the expression of that gene constitute extrinsic noise (e.g., iron manipulation in Fig. 3c). That memory could be prolonged beyond G4 by tweaking intrinsic or extrinsic components, possibly due to an increase in signal-to-noise ratio^76^, can be interpreted as a form of conditioning^77–79^. Additionally, a continuous loss of the Fe-S proteins due to noisy dilution of these proteins during cell division^80^ and damage during aerobic respiration^81, 82^ may further contribute to phenotype switching.

The incorporation of a delay in the time of transition to another state was necessary to reproduce the experimentally obtained fractions of different cell states (Figs. 6b-d). Time delay often arises from operation of a feedback^83^, or a slow incorporation of the input signal^84, 85^, or a slow response time^84, 86^. The differences in the delays specific to different states (for example, it took 3 generations for XS/M cells to switch vs 4 for L cells; Fig. 6b-d) can be due to differences in feedback time or the number of steps involved in switching. Although the *fur* regulon operates on a negative feedback loop (Fig. 3f)^87^, the expression of *fur* itself is under positive regulation of the upstream regulators Crp^88^, OxyR^89^, and SoxS^90^. This complex genetic circuit could have different feedback times depending on the cell states. For example, XS G0 mothers and their daughters would be in a Fur-repressed state till G3-4 when their iron pool goes below the threshold, so they switch to the Fur-derepressed M state. This would initiate synthesis of iron import proteins. M cells could stay derepressed for another 3-4 generations, where they accumulate enough iron to switch to the L state. This timescale is consistent with previous calculations for *fur* regulon^87^. In contrast, the switch of L back to M would require active degradation of the newly synthesized iron import proteins and mRNAs, which could take longer.

### Evolutionary significance of using iron as memory

Iron metabolism is critical for life^91^ and for its evolution^92, 93^. Both iron-limiting and iron-rich conditions can be beneficial^35, 94^ or harmful^95, 96^. So, cells should not prolong their stay in either environment. This constant switching between different iron memory states can act as a bet-hedging strategy, common all organisms^97^. Changes in iron metabolism have been reported to affect mutation rates^62^, which can directly affect natural selection^98^. Iron levels can also influence the rate of antibiotic resistance^99^, host-pathogen interaction^100–102^, composition of the gut microbiome^19^, and various other stresses in both clinical and natural settings^103^. Given the importance of iron in diverse environments^51^, iron memory is expected to be widespread in nature and to impact other physiologies. In this study we tested two other pathways influenced by iron: entering the biofilm state and resistance to antibiotics. Swarming and biofilm formation are two opposite lifestyles. While low iron is a signal for swarming, high iron was observed to be a signal for biofilms (Fig. 5a). Low iron was also positively correlated with increased survival to antibiotics (Fig. 5b, c; red circles), likely because less cellular damage is expected when iron levels are low. Antibiotics trigger production of ROS^64^, and high iron levels would increase ROS, increasing lethality^35^. A multigenerational iron memory would improve the survival chances of at least some individuals within the population under antibiotic stress. Thus, iron memory is not restricted to swarming decisions alone, and may impact additional physiologies not tested here.

Why might iron pools encode memory? Given the central role of iron in cellular metabolism, an iron-based memory offers the advantage of providing a hub connecting various stress responses. For example, swarm cells are subject to surface stress^30^ while at the same time exhibiting adaptation to antibiotic stress^44^. Changes in iron outside are not necessary to change iron inside, given that multiple other stimuli can enforce this memory.

### Limitations

1) Although we have provided a molecular framework by which iron memory operates, investigating the exact molecular basis of this memory is beyond the scope of this study. 2) An agent-based simulation is necessary to recapture the details of switching events with a feedback loop. 3) Data for the in-between generations (G2-G3, G5-G6) would have been ideal to collect, but lack of this information does not detract from the findings.

## Methods

### Strains, plasmids, media, and genetics

All bacterial strains, plasmids, and oligonucleotides are listed in Supplementary Table 1. The strains were propagated in LB broth (10 g/L tryptone, 5 g/L yeast extract, 5 g/L NaCl) or in 1.6% Bacto agar plates with the following antibiotics for marker selection as applicable – Kan (Kanamycin) 25 μg/ml, Amp (Ampicillin) 100 μg/ml, Cam (Chloramphenicol) 30 μg/ml.

*fepA* and *fur* genes were deleted in *E. coli* MG1655 by transferring the Kan^R^ deletion marker from appropriate donors strains in the Keio collection^104^ using P1 transduction (P1vir)^105^, and the deletions were confirmed by *fepA*-F/*fepA*-R and *fur*-F/*fur*R oligo combinations respectively.

### Collection of S cells

S cells were collected by flushing the actively moving edge of a 12h old swarm colony with 2 ml LB broth. After a quick spin of 1000g for 2 sec to remove any agar lumps that may have been flushed as well, the supernatant cell suspension was diluted to 0.5 OD_600_ for further assays.

### Imaging of plates

Swarm plates were imaged using a Canon EOS M6 Mark II camera and a Canon EF-M 28mm Macro IS STM lens with the following manual settings – ISO100, f/8, 1/160. The camera-generated jpg files were then loaded into Adobe Photoshop to perform equal amount of tone curve, exposure, and shadow adjustments, which was achieved by using a pre-recorded action sequence as a macro. The processed images were then exported as B&W images for further analysis.

### Colony Morphology

Hard agar LB plates (1.6% agar w/v) were dried for 8h at room temperature (RT), which is different from the normal practice of drying these plates for 24-48h before use. 100 μl of P and S cells from a 10^-5^ dilution of 0.5 OD_600_ were spread on these partially dried plates, which were incubated at 37°C for 16h and imaged. These images were processed in Adobe Photoshop as described above, then loaded in ImageJ^106^ and adjusted with a defined amount of thresholding. The circularity and diameter of the colonies was measured with the following settings of the particles: 200-infinity pixels, 0.3-1.00 circularity. ImageJ calculates circularity from the following equation: circularity = 4π(area/perimeter_2_); a perfect circle has a circularity of 1. The diameter was measured as the maximum distance between any two given points on the perimeter of the colony.

### Dilution method

The overall dilution-to-extinction method^46^ used in this study is summarized in Extended Data Fig. 1a. Instead of FACS-based sorting, dilution was chosen as the primary method of single cell isolation because of its low cost and the relative ease of inoculating a large number of swarm plates immediately after final dilution. First, the OD_600_ of the starting cell suspension – either P or S – was adjusted to 0.5. Then, 100 μl of cell suspension was serially diluted by adding 900 μl of LB medium in each step. Any recalibration or change in pipettes (or change in personnel) was accompanied by redoing the standardization steps. The accuracy of dilution was always confirmed by CFU counts. Ascertaining that the fraction of wells with growth in a 96-well plate followed a Poisson distribution, was used as an additional confirmation (Extended Data Figure 1b).

### SCI swarm assays

SCI swarm plates were prepared with 0.45% Eiken agar (Eiken Chem. Co. Japan) and 0.5% glucose. Swarm plates were dried for 8h at RT before use. 4 μL of the cell suspension was inoculated at the center of each plate and incubated at 30°C for 30h. The maximum diameter of each swarm was physically measured with a ruler. For those swarms with low radial symmetry, i.e., with no apparent max diameter upon visual inspection, three measurements with a ruler were made at different degrees of rotation to the center of the plate, and the highest value chosen as the max diameter. The highest value obtained was recorded to be the diameter.

### Zone Adjustment

Since different inoculum sizes showed large variability in the lag period before initiating swarming, a time-point based comparison of diameters across groups would not be ideal because the mean values will vary greatly. To alleviate this problem, a zone-adjustment was performed to classify data from each group into specific zones. Three arbitrary zones were assigned – central (C, 0-20 mm), medial (M, 21-60 mm), outer (O, 61-85 mm). When at least 5 members of a given group entered in a zone, swarming diameters of all the members within that same group were classified as belonging to that zone. Taking different lag durations in each group into account, 18h, 12h, and 10h data of SCIs, 100, and 10000 groups, respectively, were classified as C. For ‘M’, these values were 24h, 18h, and 12h, and for ‘O’ they were 30h, 24h, and 24h. The frequency distributions of swarm diameters for each group were then calculated keeping a swarm diameter bin size of 5 mm.

### ’Generation’ Swarm Assay

The ideal time points for daughter cell isolation from a specific generation was first standardized as summarized in Extended Data Fig. 2a-c. A suspension of 0.5 cell/ 4 μl was prepared as described in the ‘Dilution method’ section and used to inoculate 96-well plates pre-filled with 196 μl of LB medium. The plates were covered with lids, taped on the sides with parafilm, and then incubated at 37°C with shaking at 200 RPM. The number of cells per well were estimated by CFU counts at different times (see Extended Data Fig. 2a). Expectedly, some wells did not receive any cells (Extended Data Fig. 2b). Once the timepoints for specific generations were identified, the 96-well plates were inoculated and incubated as before, except, instead of CFU counts, the cells from G0, G4, and G7 were spotted on swarm plates (0.5 cell per plate) as summarized in Extended Data Fig.2d-f.

### Environmental and Genetic Perturbations

P cells were grown till 0.5 OD_600_ in 2 ml media under various environmental conditions. M9 medium (1x M9 salts, 2mM MgSO_4_, 0.1mM CaCl_2_, 0.4% glucose, 0.2% casamino acid) was used as non-rich medium, 0.1% or 10% NaCl was used for salt stress, acidic conditions were created by adjusting the pH of LB media to 6.0 or 8.0 by using 0.1 M HCl or NaOH respectively, oxygen availability was reduced by either maintaining culture tubes on bench (0 RPM) or with low shaking (50 RPM), and 0.01% or 0.05% Eiken agar was added to the LB liquid media to mimic the presence of agar in swarm plates. Cloned genes from the pCA24N plasmid-based ASKA library collection^107^, were induced with 0.1 mM IPTG. The *flhDC* clone was overexpressed from pBAD33 with 0.1% arabinose induction. To create iron-starved or iron-replete conditions, P cells were grown till 0.5 OD_600_ in 2 ml LB with 1 mM deferoxamine mesylate (DFO, Sigma) or 1 mM FeCl_3_ respectively. G0, G4, and G7 cells were collected as described in the ‘’Generation’ Swarm Assay’ section For G12, cells were collected at 300 minutes after inoculation in wells.

### Iron Biosensor

A Gibson assembly protocol^108^ was used to create the iron biosensor. First, the P*_fepA_* from WT genome, sfGFP from pMAZ plasmid, and the pTrc backbone excluding P_trc_ promoter were PCR-amplified using primer combinations *fepA*-ins-F/ *fepA*-ins-R, sfGFP-ins-F/ sfGFP-ins-R, and pTrc-F/pTrc-R, respectively. For Gibson assembly of P*_fepA_*sfGFP, all three amplicons were incubated with 2.67 μl 5X ISO buffer, 0.05 of U T5 exonuclease (NEB), 0.34 U of Phusion polymerase (NEB), 53.34 U of Taq ligase (NEB) in a 20 μl reaction. The overlapping regions at the junction of P*_fepA_* and sfGFP fragments, and complete sequence of the clones were checked using primers *fepA*-check-F/sfGFP-check-R and Seq-check-F/Seq-check-R, respectively. Separately, control plasmid sfGFP, driven by P_trc_, was also constructed using Gibson assembly with fragments of sfGFP and pTrc plasmid backbone including P_trc_.

### Microscopy

All cells were visualized using a light microscope with fluorescent channels (BX53F; Olympus, Tokyo, Japan). sfGFP was used to record fluorescence intensities from a GFP filter (Ex, 460– 480 nm; Em 495–540 nm) keeping a constant camera exposure time of 100 ms. The outputs from the 100X objective were recorded on XM10 CCD sensor (Olympus, Tokyo, Japan) using cellSens software (v1.6).

### FACS

WT and Δ*fepA* P cells harboring various reporter plasmid constructs, were grown till 0.5 OD_600_ in LB at 37°C, then collected by centrifugation, and resuspended in PBS. Cells were sorted on a BD Aria Fusion flow cytometer with BD FACSDiva (v.9.0.1) software (BD Biosciences) using a 488 nm excitation laser (530/30 band pass) and a gating strategy as illustrated in Extended Data Fig. 5a (see also Fig. 4c). The FACS sorter cannot deposit mother cells directly on a plate, so each sorted cell was deposited in one well of a 96-well plate prefilled with 20 μl LB, and then used for follow-up experiments as illustrated in Fig. 4d.

#### Swarm

For G0 mother data, each sorted cell was immediately collected from its well and spotted on a swarm plate. For G4-G7 data, 180 μl of LB was added to each sorted cell in its well. The 96-well plate was sealed and incubated as before to get G4 and G7 daughters.

#### Biofilm

A standard microplate biofilm assay^61^ was performed with minor modifications. Briefly, 80 μl of LB was added to each sorted cell in its well. The 96-well plates were then sealed with parafilm and incubated in 37℃ for 24h. The plates were washed three times by dipping them into a bucket filled with deionized water pouring out the residual water. Each well was then filled with 125μl of 0.1% crystal violet and incubated for 15 min in RT. The plates were again washed with water and then incubated in 42°C without a lid for 1.5h. 125μl of 30% acetic acid was added to each well and plates incubated at RT for 15min. The solubilized crystal violet in each well was transferred to a new well of a fresh 96-well plate. The A_550_ reading was used as a measure of biofilm formation with 30% acetic acid serving as a blank control.

#### Antibiotic survival

To measure the survival of sorted cell in sub-MIC levels or antibiotic, each sorted cell was incubated with 2 μg/ml Kan or 1.5 μg/ml Cam in a final volume of 100 μl LB. The plates were sealed as before and incubated at 37°C with 200 RPM shaking for 4h and the growth was measured as OD_600_.

### Swim assays

Fischer agar plates (0.3%) were used for swim motility assays by stabbing 4 μL of the cell suspension at the center, followed by incubation at 37°C.

### Statistics and visualization

Statistical analyses were performed using Prizm v9 (Graphpad), Python (ggplot and numpy packages) using Jupyter, and Microsoft Excel. The data was visualized in Prizm and Python.

#### PCA

To identify environmental or genetic conditions that alter the swarming heterogeneity of G0 mothers, a PCA (Principal Component Analysis) was performed on the swarm diameter data shown in Fig. 3a. The calculated mean (*m*), SD (*σ*), and the coefficient of variation (*η*) of each sample was used as three variables for PCA^48^. The resulting PC1 (60.06%) could separate *fepA* and *fur* samples from others. The PCA biplot indicated that *m* and *η* influenced PC1 the most, while *σ* and *η* influenced PC2 the most (Supplementary Fig. 3b).

#### Growth and motility clustering

The differences in growth and motility profiles of swarm colonies were measured and compared as illustrated in Extended Data Fig 3a. The images of swarm plates were processed on ImageJ with equal thresholding to measure the total area (*α*). The mean pixel intensity (*m*) was measured using the IntDen function of ImageJ. The growth variable (*V_G_*), i.e., the total pixel intensity of a swarm colony, was calculated as *m* × *α*. The motility variable (V_M_), i.e., the maximum diameter of a swarm colony (*l*) was measured using a ruler. The *V_G_* and *V_M_* variables were min-max normalized within each cohort and clustered using KMeans package of sklearn module in Python.

#### Noise calculation and F-test

The coefficient of variation (*c*) was calculated to measure noise for each sample group as *c* = _*σm*_

where *σ* and *m* denote standard deviation and mean respectively. The significance of differences among the *c* values across samples were measured by calculating p values from F-test^109^ involving the following equation

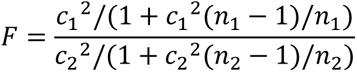

where *n* is sample size, *α* = 0.05; 1 and 2 represent groups 1 and 2 respectively.

#### Dendrogram

To illustrate how a cell type can generate other cell types across generations, a dendrogram was created using the RAWGraphs package^110^. The fractions of different types of daughters after each generation, obtained from our mathematical modeling data, were used to calculate the approximate number of each type.

### Mathematical Modeling

Classifying cells as poor (< 35 mm, XS), moderate (35-65 mm, M), and efficient swarmers (> 65 mm, L), we analyzed data from single mothers to quantify the population phenotypic heterogeneity with approximately 52%XS, 35% M, and 13% L swarmers. Tracking the state of individual cells in an expanding lineage reveals that the state of the initial single cell is maintained for approximately 4 generations (Fig. 2). However, this memory is transient and by 7 generations the lineage recreates the original population heterogeneity. We capture this transient memory using the following differential equation model:

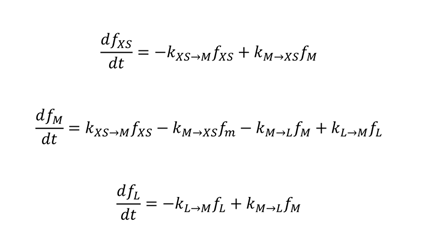

where *f_XS_*, *f*_*M*_ and *f*_*L*_ represent the fraction of cells within a single-cell derived colony at time *t* that are XS, M, and L swarmers, respectively. Here, time is normalized to the cell-cycle time so *t* = 1 corresponds to one generation. The four parameters *k*_*XS→M*_, *k*_*M*→*L*_, *k*_*M→L*_, *k*_*L*→*M*_ are the kinetic rates of transitions between the different swarming states with *k*_*XS*→*M*_ being the transition rate from XS to M swarmer, and so on. To fit the model to data we further consider a delay, where the initial cell retains its state for a certain time before starting to switch. This can be incorporated into the model by considering *k*_*XS*→*M*_ = 0 *if* *t* < τ_*XS*_ (when the initial cell is XS), *k*_*M*→*XS*_ = *k*_*M*→*L*_ = 0 *if* *t* < *τ*_*M*_ (when the initial cell is M) and *k*_*L→M*_ = 0 *if* *t* < *k*_*L*_ (when the initial cell is L). Outside these delays, these rates take constant values that are chosen such that *k*_*L→M*_ = 1.49 *k*_*XS*→*m*_ and *k*_*M→L*_ = 0.37 *k*_*L→M*_ to ensure that after some time the steady-state fractions converge to 52% XL, 35% M, and 13% L swarmers. To estimate the parameters *k*_*L*→*L*_, *k*_*L*→*M*_, *k*_ML_, *k*_*M*_, *k*_*L*_, we perform a least-squares fit between the model prediction at *t* = 4 and data at G4 (Fig. 2), and model prediction at *t* = 7 and the observed population heterogeneity of 52% XS, 35% M, and 13% L. The least-square fitting was done in Microsoft Excel using the Solver toolbox using the GRC Nonlinear solving method. The results of fitting are shown in Fig. 6b-d where the model captures the data at G4 and also recapitulate the population heterogeneity by the seventh generation. Based on the fitting we estimate the delays to be *τ*_*XS*_, *τ*_*M*_ ≈ 3 generations, *τ*_*L*_ ≈ 4 generations *k*_*XS*→*M*_ ≈ 0.4, *k*_*L*→*M*_ = 1.8 per unit time.

## Acknowledgements

We thank the following colleagues at UT Austin for their help – Alexandra Mey for suggestions with the iron biosensor design, Richard Salinas for assisting with FACS, and Yunesahng Hwang for assisting with biofilm assays. This work was supported by Public Health Service Grant GM118085 to R.M.H.

## Author Contributions

S.B., R.M.H, & A.S. conceived the project; S.B. designed the experiments; S.B., N.B., & D.M.P. performed the experiments; A.S. performed the mathematical modeling; B.W. cloned the iron biosensor; S.B. & R.M.H analyzed and interpreted experimental data; S.B. & A.S. performed statistics; S.B., R.M.H, & A.S. wrote the paper.

## Supplementary Information

The SI file contains Supplementary Tables 1-2, Supplementary Figures 1-5, and Supplementary References.

---

**Extended Data Fig. 1.**
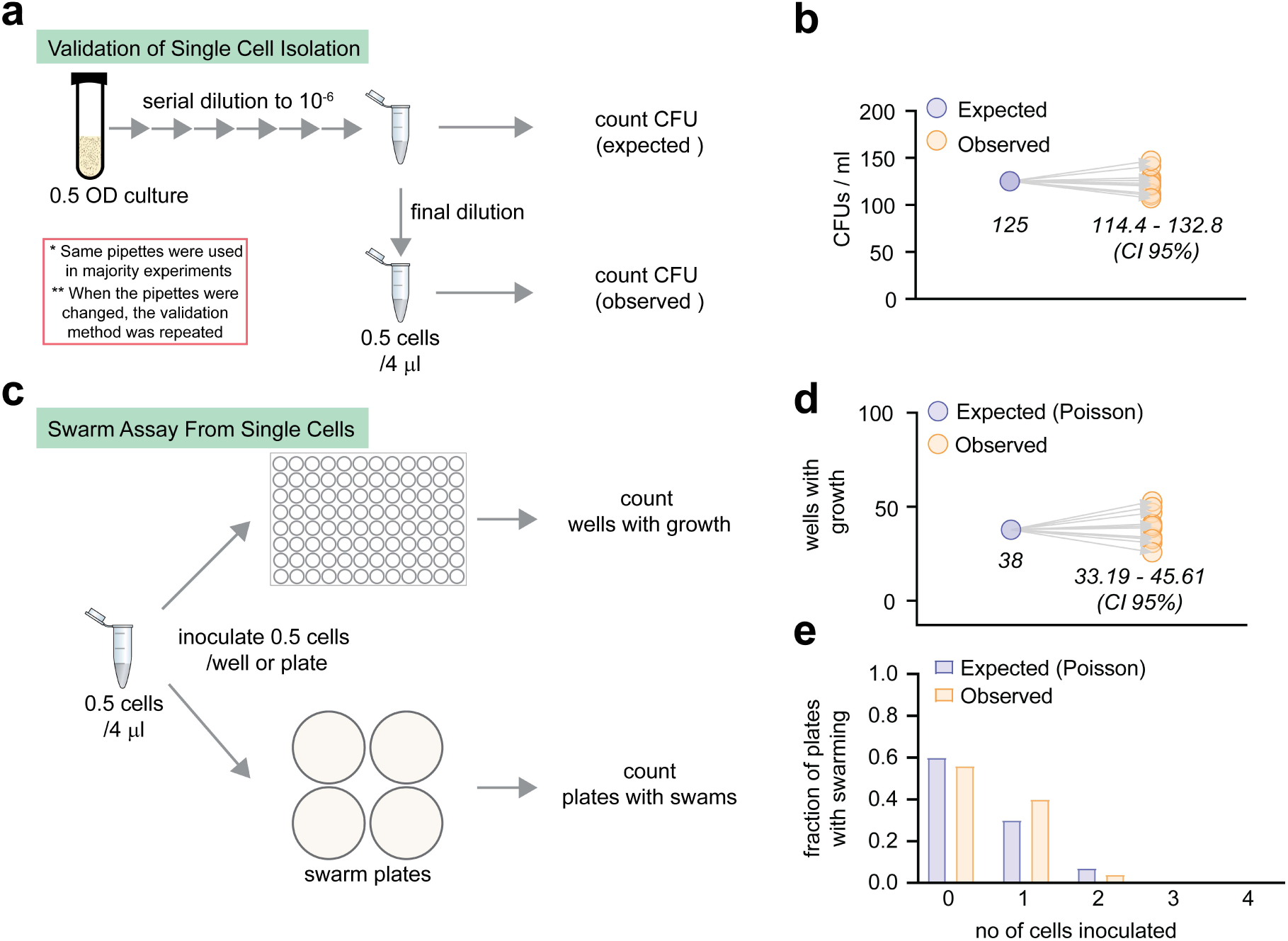
Validation of dilution-to-extinction based single cell isolation. **a**, Flow chart summarizing the single cell isolation protocol. The CFU counts of the 10^-6^ dilution (n=12) provided a mean estimate of cells per ml. This number was used to calculate the expected number of cells if diluted further. The desired dilution of 0.5 cell per 4 μl (=125 cells per ml) is obtained in the final dilution. A CFU count of this last dilution is designated ’observed’ number of cells. **b**, A comparison of the observed and expected CFUs. The confidence interval values (CI 95%) are indicated. **c**, Flow chart summarizing the steps involved in swarm assays from SCIs (Single Cell Inoculums). Since this is the first report of such methods that involve swarming, the dilution was validated in every step. **d**, The dilution of 0.5 cells/ 4 μl was further validated by seeding each well of a 96-well plate with 0.5 cells. The expected positive number of events (wells that received cells) was calculated from a Poisson distribution (μ=0.5, n=96) and compared with the observed values, i.e., number of wells with visible growth after 18h. **e**, The expected positive number of events (plates that received cells), was also calculated from a Poisson distribution (μ=0.5, n=100) and compared with the observed values, i.e., number of plates with swarm colonies after 18h.

**Extended Data Fig. 2.**
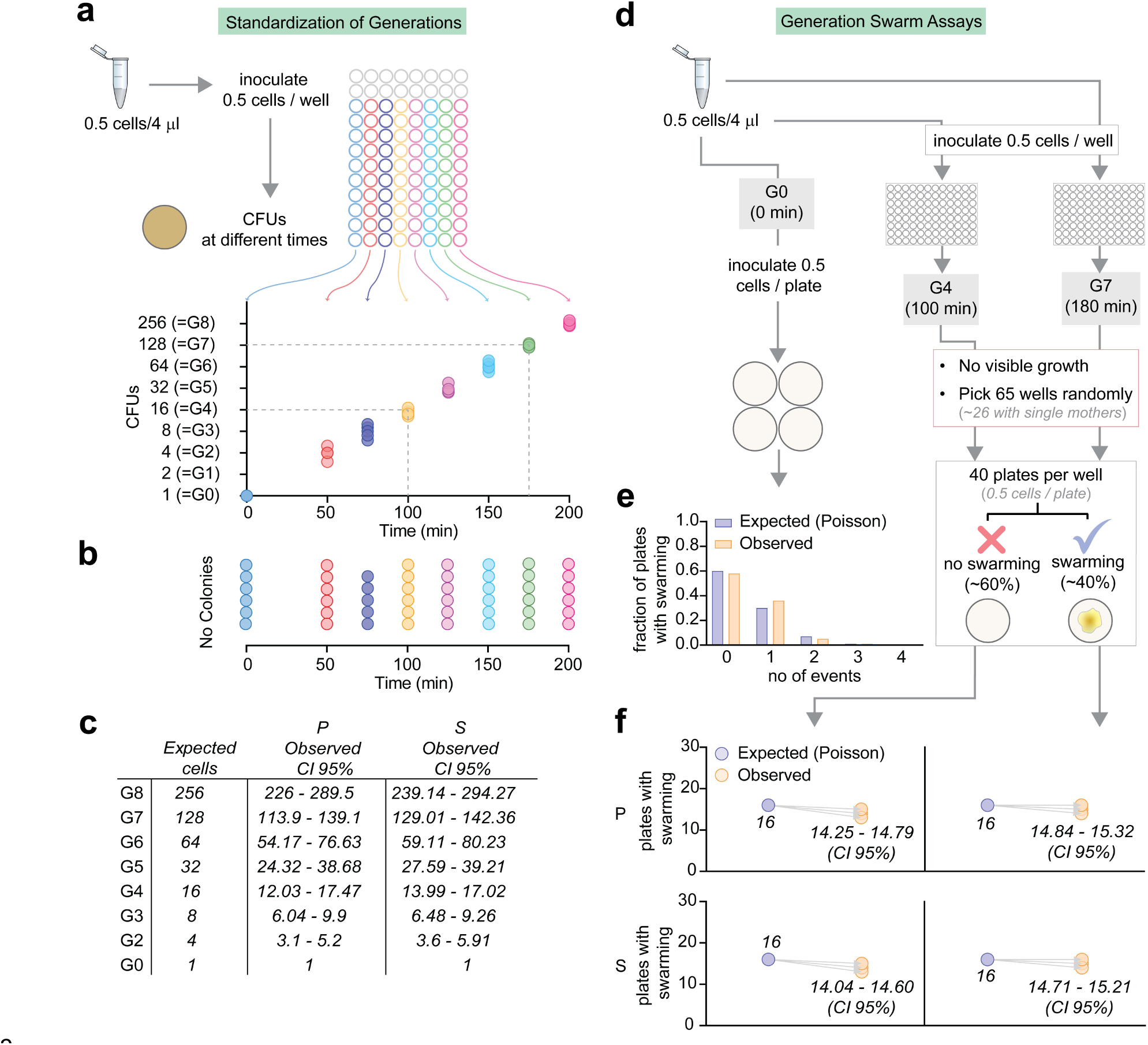
Swarm assays from single mother cells and their daughters from various generations of growth. **a-c**, Infographic showing the standardization of time points for collecting daughters from different generations. **a,** The dilution of 0.5 cells per 4 μl was used to seed single mothers in wells (colored wells). From Poisson, it is expected that only ∼40% of the wells would have received cells. CFU counts (y-axis) were performed on 10 wells at each time point (x axis) and the data from only positive events are shown in the dot plot. The expected number of cells after each generation are indicated. Only P mother data are shown here but both the P and S cells were in G4 and G7 at 100 min and 175 min respectively. **b**, The number of wells (from **a**) with no CFU (from P samples). Each dot represents a well. **c**, The confidence intervals of the observed number of cells in **a**. The data from S mothers are also shown here. **d-f**, Infographic showing the steps involved in swarm assays of SCIs collected from G4 or G7. **d**, The collected P or S cells were designated G0 immediately after the final dilution of 0.5 cells per 4 μl was made. Some G0 mothers were inoculated on swarm plates (**e**, μ=0.5, n=125 for both P and S), while other G0 mothers were seeded in wells and grown till G4 or G7. The total number of cells are very low at these time points, so no growth was visible. To get data from at least 25 mothers, 65 wells were randomly picked in each generation and processed as shown. A comparison of expected and observed positive outcomes is shown in **f**.

**Extended Data Fig. 3.**
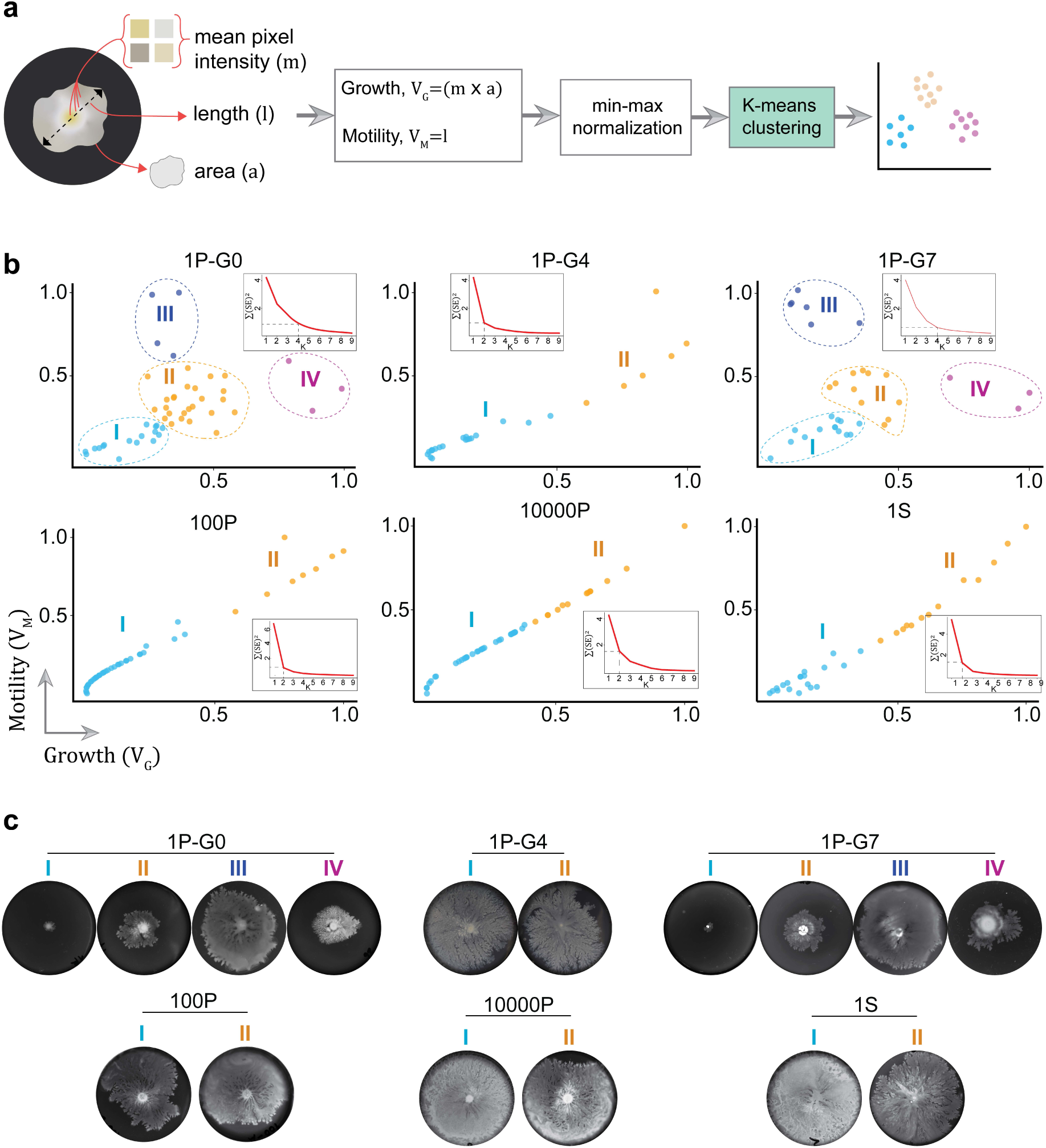
Growth and motility-based clustering of swarm data. **a**, Infographic showing the steps involved in the clustering process. The images of swarm plates were processed to measure the variables, growth (VG) and motility (VM) as the total pixel intensity and the maximum diameter of a swarm respectively (see Methods). A K-means clustering was performed within each group. **b**, The results of clustering analysis of the different datasets as labeled: 1P, planktonic mothers at G0 (n=50); 1P-G4 & 1P-G7, daughters of two random L mothers each from G4 and G7 (∼n=32 in each); 100P & 10000P, planktonic 100-and 10000-cell inoculums (n=50 in each); 1S, swarm mothers at G0 (n=49). The elbow method was used to identify the correct K value. Inset, the corresponding elbow plot showing the within-cluster sum of squares for different K values. Different clusters in each group are color-coded. **b**, Images of representative swarm plates from each cluster.

**Extended Data Fig. 4.**
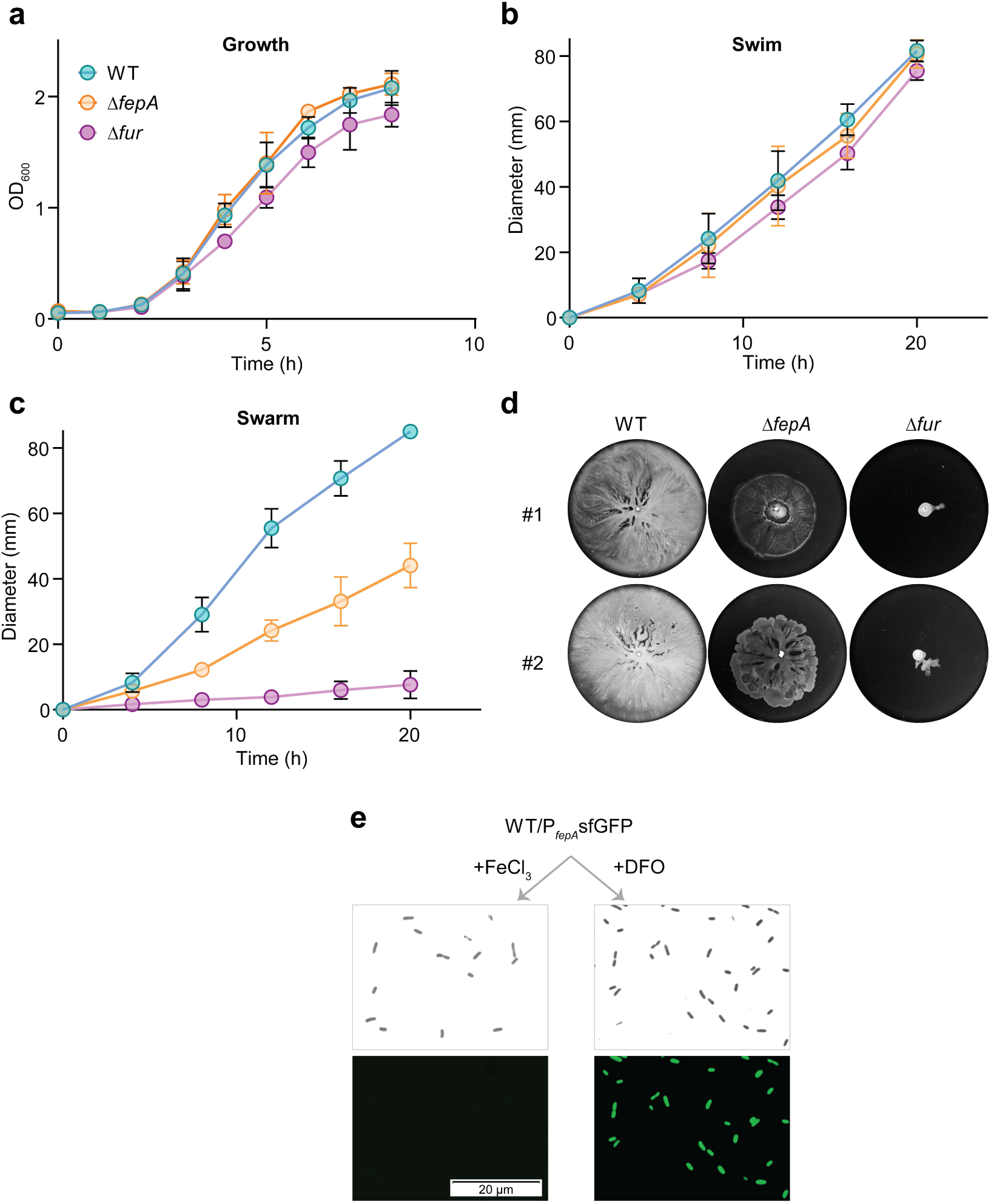
Growth & motility of iron-deficient strains. **a-c**, Time-course measurements of various phenotypes of the indicated strains (n=4): **a**, Growth curves in LB media at 37°C; **b**, Swimming motility measured (diameter) in 0.3% soft agar plates at 37°C; **c**, Swarm diameter measured on 0.5% agar at 30°C. **d**, Representative images of swarm plates of indicated strains at 20h. The numbers indicate replicates. **e**, Microscopic images of a WT strain with GFP driven from the *fepA* promoter, treated with FeCl3 or DFO.

**Extended Data Fig. 5.**
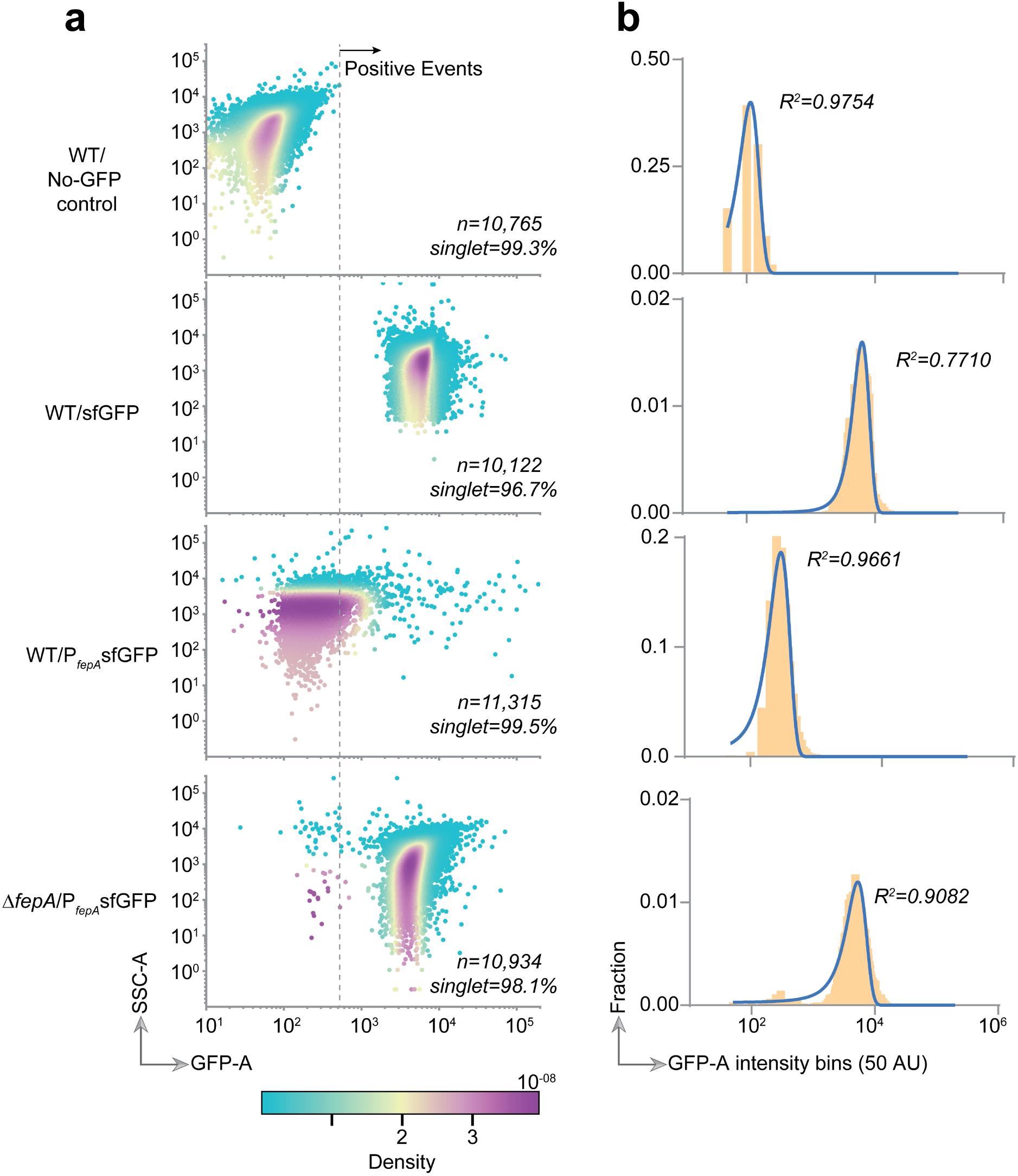
Flow cytometry of iron-biosensor strains. **a**, Flow cytometry scatter plots of singlet events, which were gated based on SSC-A (side scatter channel - area) and GFP intensity. Both x and y axes scales are biexponential. The number of events for each strain and associated singlet percentages are indicated. The heatmap is based on the kernel density estimate of the events. As a gating strategy, any singlet falling within the maximum value obtained in the GFP-A channel for the strain without any GFP (top row) was considered a negative event. The singlets above this threshold were considered positive events and sorted in individual wells of 96-well plates (see Supplementary Fig. 4 for sorting data). **b**, Frequency distributions of GFP intensity (x axis, log scale) from the corresponding strains in **a**. Note that the y-axis scales are not identical. A Gaussian fit and associated R^2^ value are shown.

