## Supplementary material for "Iron Memory in *E. coli*": Supplemenatary Information

**Affiliations:**

**Brief Contents:**

Supplementary Tables 1-2

Supplementary Figures 1 to 6

Supplementary References

**Supplementary Tables**

**Supplementary Table 1: Strains, Plasmids, and Oligonucleotides**

| **Strain/Plasmid** | **Genotype** | **Source** |
| --- | --- | --- |
| **Strains** | | |
| WT | *Escherichia coli* MG1655; F^-^ λ^-^ *rph*-1 | Lab collection^1^ |
| Δ*fepA* | MG1655 Δ*fepA::*Kan^R^ | This study |
| Δ*fur* | MG1655 Δ*fur::Kan^R^* |  |
| **Plasmids** | | |
| pCA24N | From ASKA collection, Cam^R^ | ASKA^2^ |
| *rflP* | pCA24N carrying *rflP* |  |
| *ompA* | pCA24N carrying *ompA* |  |
| *ompF* | pCA24N carrying *ompF* |  |
| *yhdC* | pCA24N carrying *yhdC* |  |
| *fepA* | pCA24N carrying *fepA* |  |
| *fur* | pCA24N carrying *fur* |  |
| *marA* | pCA24N carrying *marA* |  |
| *evgA* | pCA24N carrying *evgA* |  |
| *katG* | pCA24N carrying *katG* |  |
| *sodB* | pCA24N carrying *sodB* |  |
| *mscL* | pCA24N carrying *mscL* |  |
| *mscS* | pCA24N carrying *mscS* |  |
| *flhDC* | pBAD33 carrying *flhDC* | Partridge & Harshey^3^ |
| pMAZ-sfGFP | pMAZ-SK carrying sfGFP, Kan^R^ | Finkelstein Lab, UT Austin |
| pTrc99a | Cloning vector, P_Trc_, Amp^R^ | Lab collection^4^ |
| sfGFP | pTrc99a carrying sfGFP, P_trc_, Amp^R^ | This study |
| P*_fepA_*sfGFP | pTrc99a carrying sfGFP, P*_fepA_*, Amp^R^ |  |
| **Primers** | | |
| **Name** | **Sequence (5’🡪3’)** | **Purpose** |
| *fepA*-F | CGACCATATTCGAGACTGATG | *fepA* KO confirmation |
| *fepA*-R | AGTGATGTAGACCCATACGC | *fepA* KO confirmation |
| *fur*-F | GGTTTTCATTTAGGCGTGGCAATTC | *fur* KO confirmation |
| *fur*-R | GATAAAGTCTGGCAGGAAATACGC | *fur* KO confirmation |
| pTrc-F | GAGAAGATTTTCAGCCTGATACAG | amplify pTrc backbone |
| pTrc-R | GGATCCCTTGTCTGTAAGCGGATGCCGGGA |  |
| sfGFP-ins-F | TCGTGGCAAAAATGCAGGAATAAAACAGAATTCGGATCTTA  GCTACTAGAGAAAGAGGAG | amplify sfGFP from pMAZ-sfGFP |
| sfGFP-ins-R | TTCTGATTTAATCTGTATCAGGCTGAAAAGCTTCTCTCA  TTTGTACAGTTCATCCATACC |  |
| *fepA*-ins-F | TTGTCTGCTCCCGGCATCCGCTTACAGACAAGGGATCCGAC  TGCCACCAGCTCTCACTTC | Amplify P*_fepA_* promoter from WT |
| *fepA*-ins-R | TCTCCTCTTTCTCTAGTAGCTAAGATCCGAATTCTGTTTTAT  TCCTGCATTTTTGCCACG |  |
| *fepA*-check-F | CACTGCGTGTCTTTCAGGAT | check overlaps |
| sfGFP-check-R | GTTACCAGAGTCGGCCAAGG |  |
| Seq-check-F | CGCATCTGTGCGGTATTTCAC | sequencing |
| Seq-check-R | GAGCCTTTCGTTTTATTTGATGCC |  |

**Supplementary Table 2: Software and Algorithms**

| **Name** | **Description** | **Source** |
| --- | --- | --- |
| ImageJ v1.47 | For colony measurements | https://imagej.net/ ^5^ |
| Prism v9 | Data visualization, statistical testing | https://graphpad.com |
| matplotlib v3.5 | Python package for data visualization | https://matplotlib.org/ |
| numpy v*1.23* | Python package for data visualization | https://numpy.org/ |
| pandas v1.4 | Python package for data visualization | https://pandas.pydata.org/ |
| joypy v0.2.5 | Python package for joyplot | https://github.com/leotac/joypy |
| scipy.stats | Python package for data analysis | https://scipy.org/ |
| BDFACS Diva v9.0.1 | FACS data acquisition | https://www.bdbiosciences.com/en-us/products/software/instrument-software/bd-facsdiva-software |
| FlowJo v10.8.1 | FACS data export | https://www.flowjo.com/ |
| cellSens software v1.6 | Olympus | https://www.olympus-lifescience.com/en/software/cellsens/ |
| RAWGraphs | Dendrograms | https://www.rawgraphs.io/about^6^ |

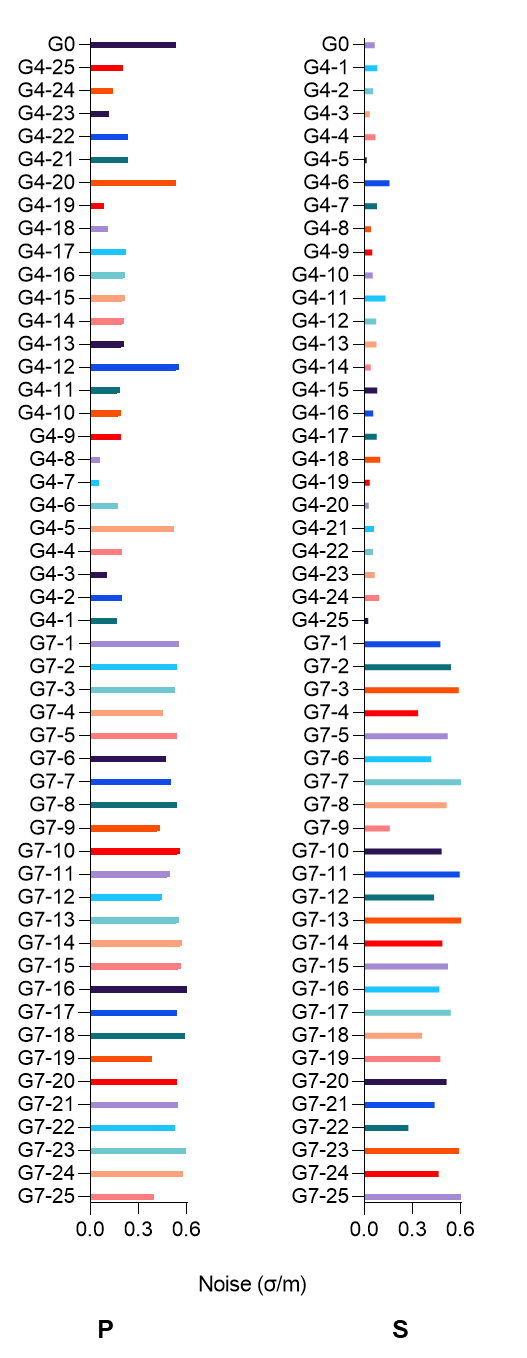
**Supplementary Fig. 1 | The noise data for Fig2.** The comparison of noise (SD/mean) for each sample from main Fig. 2.

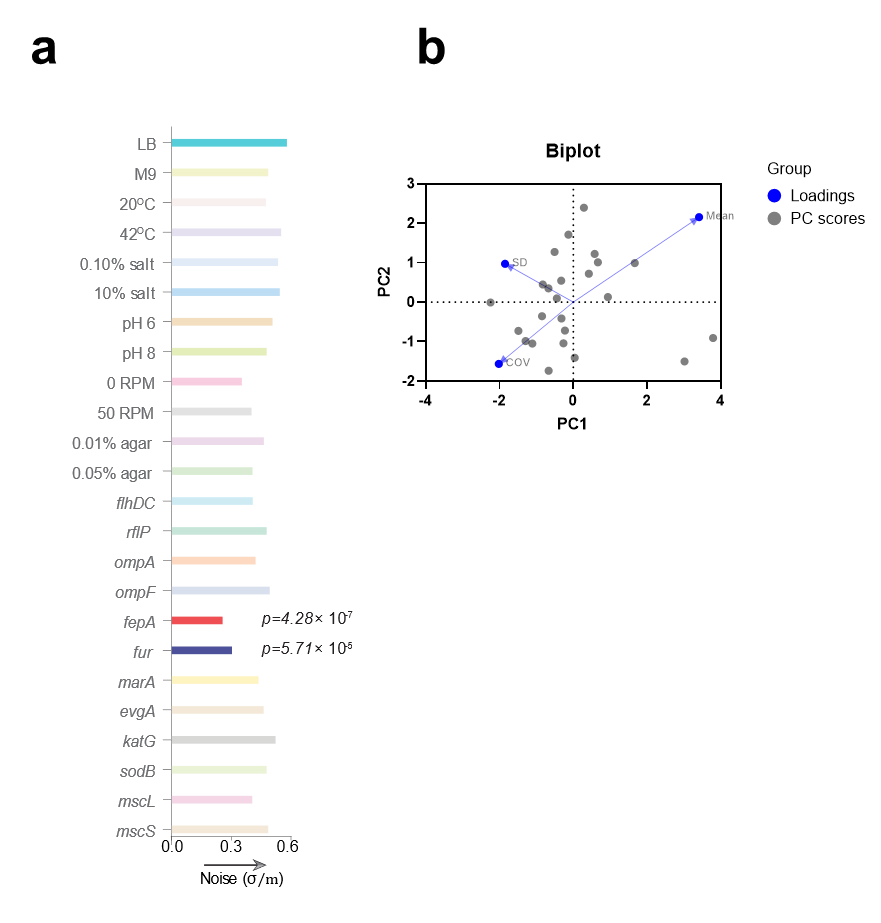

**Supplementary Fig. 2 | The noise data for Fig. 3 and PCA Biplot. a**, Comparison of noise (SD/mean) for each sample from main Fig. 3. **b**, Biplot for PCA shown in main Fig. 3b.

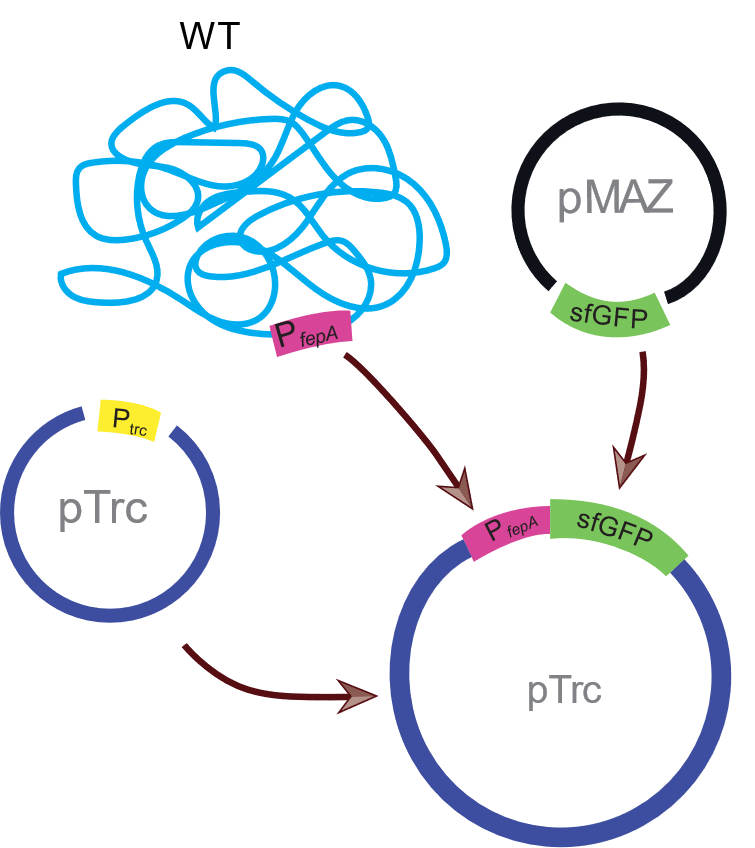

**Supplementary Fig. 3 | Cloning strategy for the iron biosensor.** Super-folder GFP (sfGFP) from pMAZ plasmid, driven by the *E. coli* *fepA* promoter (P*_fepA_*) was cloned into a pTrc vector plasmid (P*_fepA_-*sfGFP). A control plasmid with sfGFP driven from the IPTG-inducible pTrc promoter was also constructed (P_trc_-sfGFP; not shown).

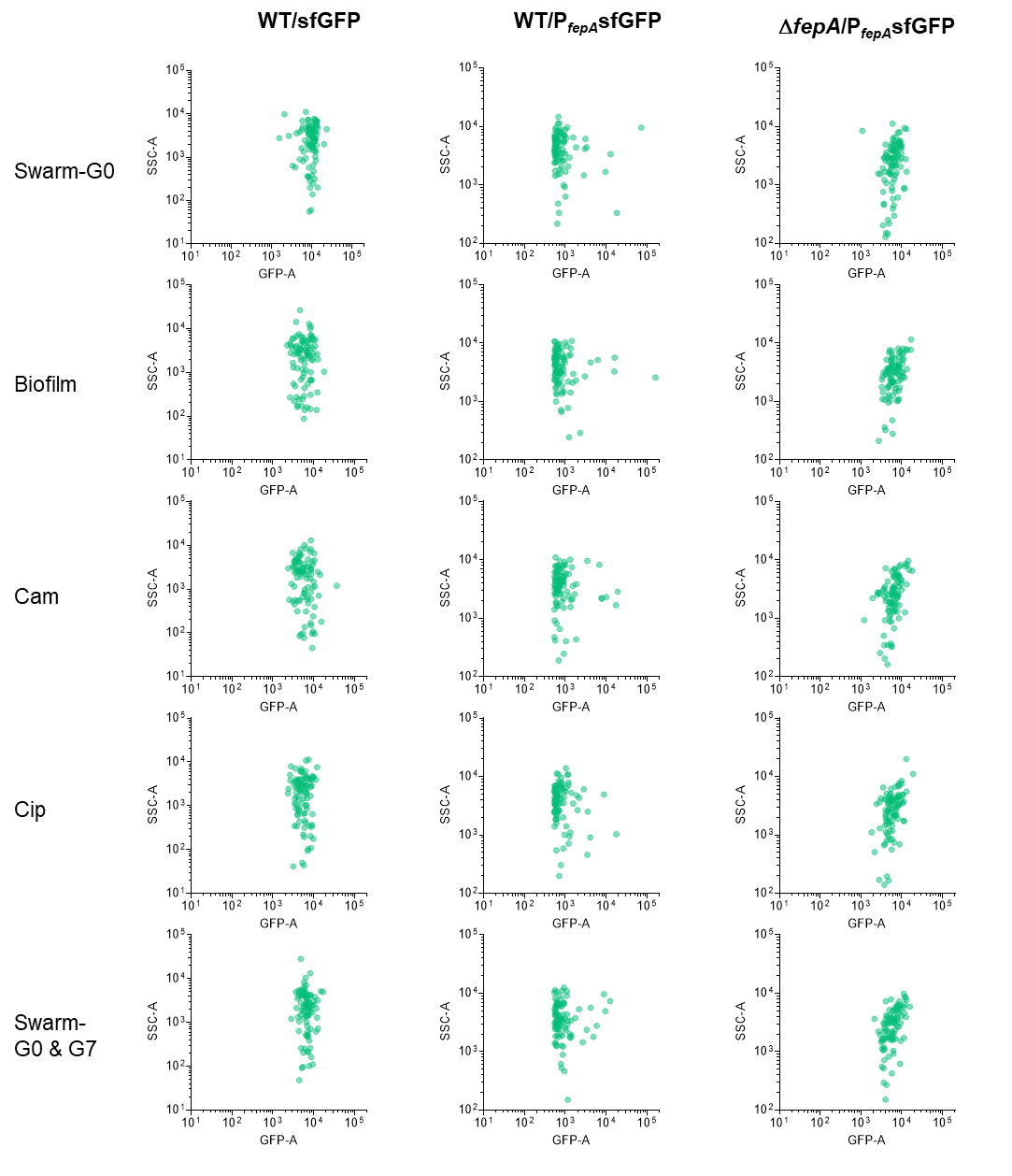

**Supplementary Fig. 4 | Flow cytometry data of the sorted cells related to Figs. 4 & 5.** The singlets were sorted based on a gating strategy as described in Extended Data Fig. 5. Each green dot represents a cell that was sorted and dropped into a well of a 96-well plate. For example, the top left panel shows data for sorted WT/sfGFP cells that were used for performing swarm assays at G0 which is shown in Fig. 4e (left).

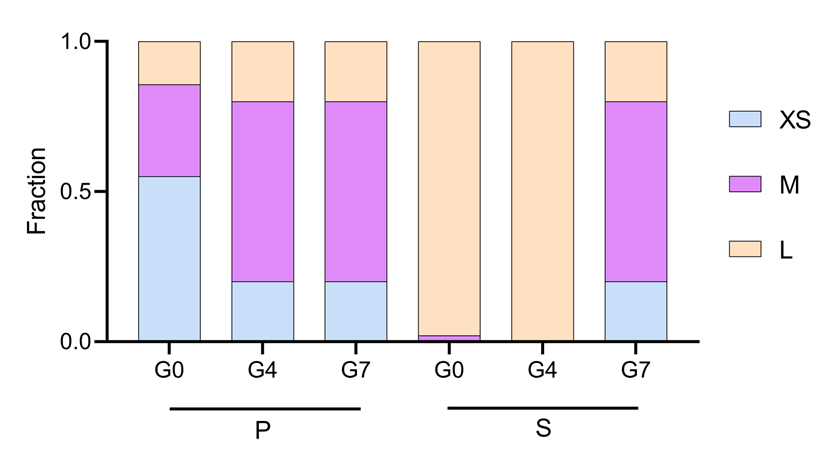

**Supplementary Fig. 5 | Fraction of different categories of swarmers from the data in Fig. 2.** See main Figure 2 for the details of clustering.

1	Guyer, M. S., Reed, R. R., Steitz, J. A. & Low, K. B. Identification of a sex-factor-affinity site in E. coli as gamma delta. *Cold Spring Harbor symposia on quantitative biology* **45 Pt 1**, 135-140, doi:10.1101/sqb.1981.045.01.022 (1981).

2	Kitagawa, M. *et al.* Complete set of ORF clones of Escherichia coli ASKA library (a complete set of E. coli K-12 ORF archive): unique resources for biological research. *DNA research : an international journal for rapid publication of reports on genes and genomes* **12**, 291-299, doi:10.1093/dnares/dsi012 (2005).

3	Partridge, J. D. & Harshey, R. M. More than motility: Salmonella flagella contribute to overriding friction and facilitating colony hydration during swarming. *Journal of bacteriology* **195**, 919-929, doi:10.1128/JB.02064-12 (2013).

4	Amann, E., Brosius, J. & Ptashne, M. Vectors bearing a hybrid trp-lac promoter useful for regulated expression of cloned genes in Escherichia coli. *Gene* **25**, 167-178, doi:10.1016/0378-1119(83)90222-6 (1983).

5	Schneider, C. A., Rasband, W. S. & Eliceiri, K. W. NIH Image to ImageJ: 25 years of image analysis. *Nature methods* **9**, 671-675, doi:10.1038/nmeth.2089 (2012).

6	Mauri, M., Elli, T., Caviglia, G., Uboldi, G. & Azzi, M. in *Proceedings of the 12th Biannual Conference on Italian SIGCHI Chapter*	Article 28 (Association for Computing Machinery, Cagliari, Italy, 2017).
